# Multifunctional CRISPR/Cas9 with engineered immunosilenced human T cell epitopes

**DOI:** 10.1101/360198

**Authors:** Shayesteh R. Ferdosi, Radwa Ewaisha, Farzaneh Moghadam, Sri Krishna, Jin G. Park, Mo R. Ebrahimkhani, Samira Kiani, Karen S. Anderson

## Abstract

The application of Cas9 for genetic and epigenetic therapies in humans raises concerns over immunogenicity of this foreign protein. We report pre-existing human CD8+ T cell immunity to *Streptococcus pyogenes* Cas9 in the majority of healthy individuals screened. In a proof-of-principle study, we demonstrate that Cas9 protein can be modified to eliminate immunodominant epitopes through targeted mutation while preserving its function and specificity.

## Introduction

The Clustered Regularly Interspaced Short Palindromic Repeat (CRISPR)/Cas9 technology has raised hopes for developing personalized gene therapies for complex diseases such as cancer as well as genetic disorders, and is currently entering clinical trials ^1,2^. The history of gene therapy has included both impressive success stories and serious immunologic adverse events ^3-8^. The expression of *Streptococcus pyogenes* Cas9 protein (SpCas9) in mice has evoked both cellular and humoral immune responses ^9,10^, which raises concerns regarding its safety and efficacy as a gene or epi-gene therapy in humans. These pre-clinical models and host immune reactions to other exogenous gene delivery systems ^11-13^ suggest that the pathogenic “non-self” origin of Cas9 may be immunogenic in humans.

Both B cell and T cell host responses specific to either the transgene or the viral components of adenoviral ^14,15^ and adeno-associated viral (AAV) ^11,12^ vectors have been detected, despite relatively low immunogenicity of AAV vectors. In the case of AAV, specific neutralizing antibodies (Abs) and T cells are frequently detected in healthy donors ^16-19^ and specific CD8+ T cells have been shown to expand following gene delivery ^18^. There has been recent progress in developing strategies to overcome this problem, such as capsid engineering and transient immunosuppression ^20-22^. The potential consequences of immune responses to expressed proteins from viral vectors or transgenes include neutralization of the gene product; destruction of the cells expressing it, leading to loss of therapeutic activity or tissue destruction; induction of immune memory that prevents re-administration; and fulminant innate inflammatory responses ^23,24^. More potent immune responses to gene therapies have been observed in humans and non-human primate models compared to mice ^8,25^.

Of the Cas9 orthologs derived from bacterial species, the SpCas9 is the best characterized. *S. pyogenes* is a ubiquitous pathogen, with an annual incidence of 700 million worldwide ^26^, but immunity to SpCas9 in humans has not been reported. Here, we sought to characterize the pre-existing immune response to SpCas9 in healthy individuals and to identify the immunodominant T cell epitopes with the aim of developing SpCas9 proteins that have diminished capacity to invoke human adaptive response.

CRISPR application for human therapies will span its use both for gene editing (through DNA double-strand breaks) or epigenetic therapies (without DNA double-strand breaks). In fact, recent reports shed light on CRISPR’s ability to activate or repress gene expression in mice ^27-29^, which opens the door to a variety of new therapeutic applications such as activating silent genes, compensating for disrupted genes, cell fate reprogramming, or silencing disrupted genes, without the concern over permanent change in DNA sequence. However, unlike the use of Cas9 for gene editing, which may only require Cas9 presence in cells for a few hours, current techniques for CRISPR-based epigenetic therapies require longer term expression of Cas9 *in vivo*, possibly for weeks and months ^28,29^, which poses the challenge of combating pre-existing immune response towards Cas9. This challenge will need to be addressed before CRISPR application for human therapies, especially for epigenetic therapies, can be fully implemented. Delivery of CRISPR *in vivo* by incorporating its expression cassette in adeno-associated virus (AAV), will most likely shape many of the initial clinical trials as AAV-based gene delivery is one of the safest and most prevalent forms of gene therapies in human. AAV will enable longer term expression of Cas9, desirable for epigenetic therapies. Therefore, unlike Cas9 delivery in the form of ribonucleoprotein complexes (which are short term), it is highly likely that CRISPR delivery through AAV and its expression within target cells will engage CD8+ T cell immunity.

## Results

We first determined whether healthy individuals have detectable IgG Abs to SpCas9. Of 143 healthy control sera screened, 70 (49.0%) had detectable Abs against *S. pyogenes* lysate using ELISA (**Fig. 1A**). This positive subset along with sera that were borderline negative for Abs to *S. pyogenes* lysate were screened for Abs against recombinant SpCas9, of which 36.6% were positive. At least 21.0% (n=30) of healthy individuals in this study had Cas9-specific Abs (**Fig. 1A**).

**Figure 1.**
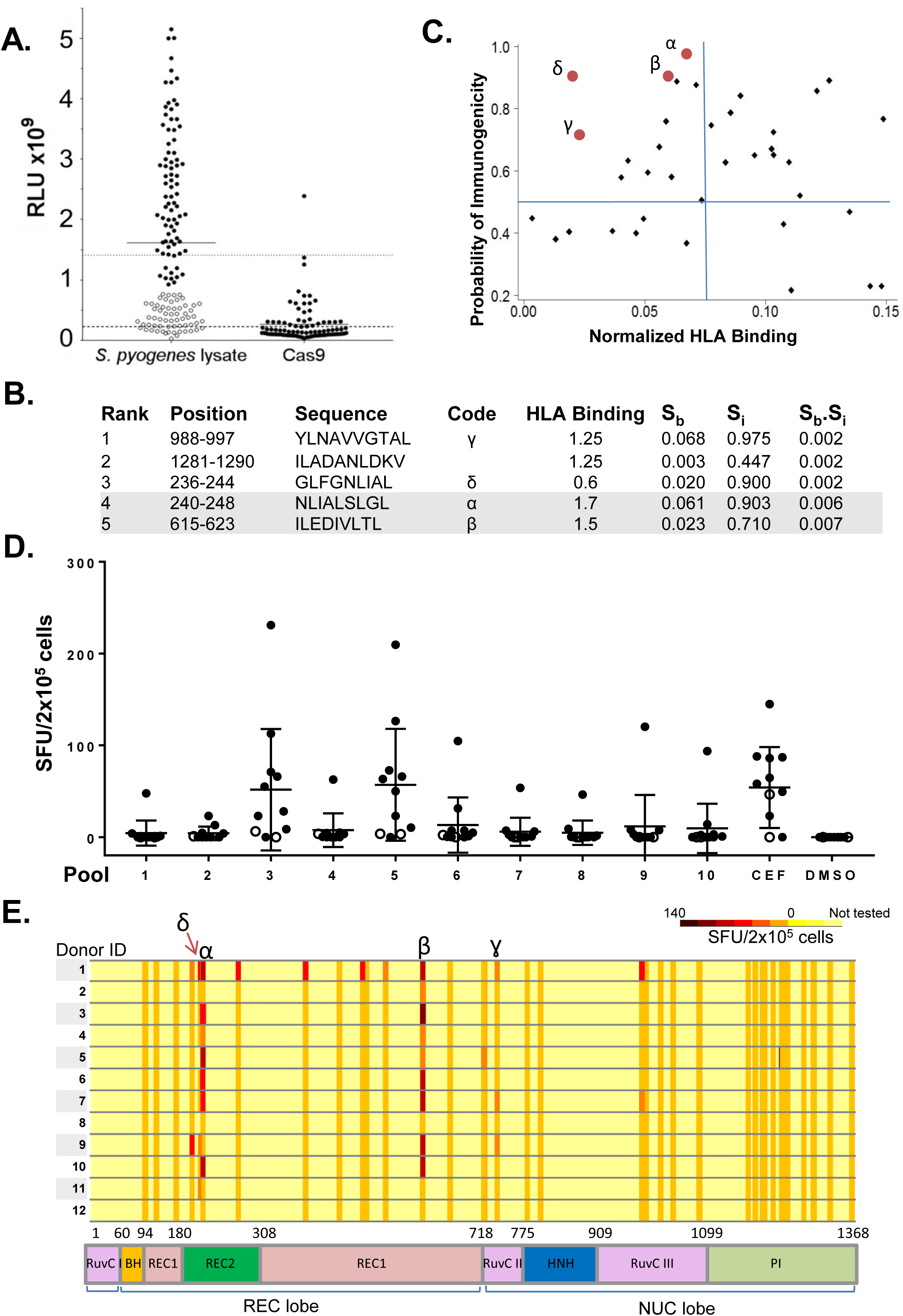
Detection of pre-existing B cell and T cell immune responses to SpCas9 in healthy donors and identification of two immunodominant T cell epitopes. **A.** Specific serum Abs were detected against *S. pyogenes* lysate in 49.0% (above the dotted line) of 143 healthy controls (left). The subset shown in black circles was screened for Abs against recombinant Cas9 protein (right), of which 36.6% (21.0% of total samples screened) were positive (above the dashed line). **B.** The top 5 predicted SpCas9 T cell epitopes and their predicted S_b_ and S_i_ scores and ranking (based on the S_b_.S_i_ value) ^30^. These top 5 peptides include the identified immunodominant (α and β; gray) and subdominant (γ and δ) epitopes that were shown to be immunogenic by IFN-γ ELISpot. **C.** Plot of S_b_ and S_i_ of predicted HLA-A^∗^02:01 epitopes for the SpCas9 protein. Red dots represent the immunodominant and subdominant epitopes. **D.** IFN-γ ELISpot assay of T cell reactivity of 12 healthy donors (the two non-HLA-A^∗^02:01 are shown as open circles) to 38 predicted epitopes grouped in 10 pools, CEF (positive control), and DMSO (negative control). Peptides α and δ were in pool 5 while β and γ were in pool 3. **E.** IFN-γ ELISpot reactivity of healthy donor T cells (n=12) to epitopes across the different domains of the Cas9 protein. Donors 1-10 were HLA-A^∗^02:01, while 11 and 12 were not. Peptides α and δ overlap in 5 amino acid residues. S_b_, normalized binding score; S_i_, normalized immunogenicity score.

Whether Cas9-specific antibodies impact the efficacy or safety of CRISPR application in human remains to be seen. However, cellular immunity is expected to have a more significant impact given the current CRISPR/Cas9 delivery approaches. The gene encoding Cas9 is usually delivered by a vector such as viral vectors to target cells and for intracellular expression, which could evoke a cellular immune response. We thus focused on investigating the T cell immune response against SpCas9. We predicted HLA-A^∗^02:01-restricted T cell epitopes derived from SpCas9 using a model that uses both MHC binding affinity and biochemical properties of immunogenicity ^30^ (**Supplementary Table 1**; the top 5 are shown in **Fig. 1B**). This model incorporates T cell receptor contact residue hydrophobicity and HLA binding prediction, which enhances the efficiency of epitope identification, as we previously reported ^30^. We plotted the calculated normalized binding (S_b_) and immunogenicity (S_i_) scores for each peptide (**Fig. 1C**) to predict the more immunogenic epitopes, which are expected to have both high HLA binding (low S_b_) and more hydrophobicity (high S_i_). We chose HLA-A^∗^02:01 because it is the most common HLA type in European/North American Caucasians. We also predicted MHC class II binding epitopes for the SpCas9 protein to HLA-DRB1 (10 alleles), HLA-DQ (5 alleles), and HLA-DP (8 alleles) using the IEDB analysis tool (**Supplementary Table 2**).

We then investigated whether peripheral blood mononuclear cells (PBMCs) derived from healthy individuals had measurable T cell reactivity against the predicted SpCas9 MHC class I epitopes. We synthesized 38 peptides (**Supplementary Table 1**) and grouped them into 10 pools of 3-4 peptides each. We measured peptide-specific T cell immunity using IFN-γ secretion ELISpot assays with PBMCs derived from 12 healthy individuals (HLA-A^∗^02:01, n=10; non-HLA-A^∗^02:01, n=2) and identified immunoreactive epitopes within pools 3 or 5 in 83.0% of the donors tested (90% of the HLA-A^∗^02:01 donors; **Fig. 1D**). The seven individual peptides from pools 3 and 5 were evaluated by IFN-γ ELISpot and the dominant immunogenic epitopes were SpCas9_240-248 and SpCas9_615-623, designated peptides α and β, from pools 5 and 3, respectively. The subdominant epitopes were found to be γ and δ from pools 3 and 5, respectively. Both peptides α and β are located in the REC lobe of the Cas9 protein (**Fig. 1E)** that binds the sgRNA and the target DNA heteroduplex ^31^. The individual peptides within pools that were positive for any donor were evaluated for this donor by IFN-γ ELISpot. The immunoreactivity and position of the 38 predicted peptides (a few of which are overlapping) within the Cas9 protein are shown in **Fig. 1E**.

Peptides α and β are shown as red dots on the epitope prediction plot (**Fig. 1C**) and their sequences and predicted ranking are shown in **Fig. 1B** and **Supplementary Table 1**. As predicted, these peptides had low S_b_ and high S_i_ values. Both the immunodominant (α and β) and subdominant (γ and δ) T cell epitopes identified by IFN-γ ELISpot were within the top 5 most immunogenic epitopes predicted by our immunogenicity model ^30^. Their ranking as predicted by the consensus method hosted on the IEDB server using default settings was 14, 5, 18, and 4, respectively. For MHC class II, epitope α is predicted to be a top binder to HLA-DRB1^∗^01:02 and epitope β a top binder to HLA-DPA1^∗^01:03 and DPB1^∗^02:01 (**Supplementary Table 2**). Sequence similarity of peptides α and β to amino acid sequences in known proteins was investigated using Protein BLAST and the IEDB epitope database ^32^. This was done to investigate whether there is any chance that the T cell immune response that we are detecting in healthy individuals could be due to previous exposure to another protein of similar sequence. A peptide was considered ‘similar’ to α or β if no more than 2 of 9 amino acid residues (that are not the second or ninth) were not matching (78% similarity). None of these two peptides resembled known epitopes in the IEDB database, but similarity to other Cas9 orthologs and other bacterial proteins was detected (**Supplementary Tables 3** and **4**). Epitope β has sequence similarity to a peptide derived from the *Neisseria meningitidis* peptide chain release factor 2 protein (ILEDIVLTL versus ILE**<u>G</u>**IVLTL). Antigen-specific T cells were expanded for 18 days *in vitro* by coculturing healthy donor PBMCs with peptide β-pulsed autologous antigen presenting cells (APCs). Cas9-specific CD8+ T cell responses were assessed by flow cytometry. CD8+ T cells specific for the HLA-A^∗^0201/β pentamer were detected after stimulation (3.09%; **Fig. 2A**).

**Figure 2.**
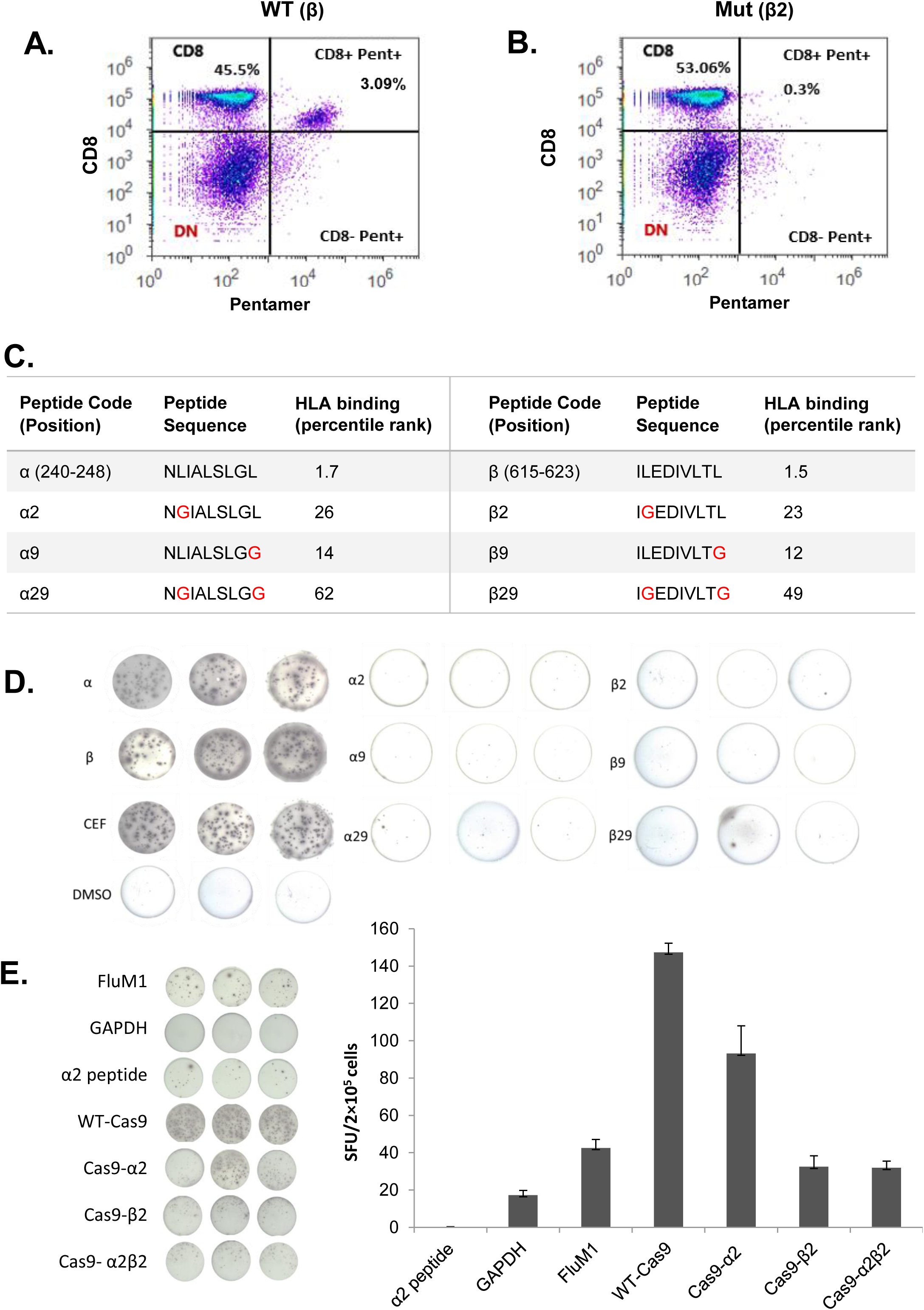
SpCas9 immunodominant epitope-specific CD8+ T cell recognition is abolished after anchor residue mutation. **A.** Epitope β-specific CD8+ T cell response detected using β-specific pentamer in PBMCs stimulated with peptide p-pulsed antigen presenting cells. **B.** The percentage of CD8+ pentamerβ+ T cells was reduced to 0.3% when healthy donor B cell APCs were pulsed with the mutated peptide β2. **C.** Positions, sequences and IEDB HLA binding percentile rank of epitopes α and β before and after mutation of the anchor (2^nd^ and/or 9^th^) residues. S_b_, normalized binding score; S_i_, normalized immunogenicity score. **D.** IFN-γ ELISpot assay in triplicate wells comparing T cell reactivity to wild type or mutated epitopes α and β. These results are representative of 12 donors and two independent replicates. **E.** IFN-γ ELISpot comparing T cell reactivity to APCs expressing WT or modified Cas9 proteins. APCs expressing FluM1 were used as a positive control. APCs expressing GAPDH or spiked with peptide α2 were used as negative controls. Data represent mean +/- SEM of 5 replicates (right).

We next hypothesized that mutation of the MHC-binding anchor residues of the identified immunogenic epitopes would abolish specific T cell recognition (**Fig. 2A**). The epitope anchor residues (2^nd^ and 9^th^) are not only necessary for peptide binding to the MHC groove, but are also crucial for recognition by the T cell receptor ^30^. The percentage of CD8+ β pentamer+ T cells decreased to 0.3% when APCs were pulsed with the mutated peptide (β2; **Fig. 2B**) compared with 3.09% with the wild type peptide (β; **Fig. 2A**). We then examined the reactivity of healthy donor T cells to modified peptides α or β with mutations in residues 2, 9, or both (sequences are shown in **Fig. 2C**) using IFN-γ ELISpot assay. The epitope-specific T cell reactivity was markedly reduced with the mutant peptides (**Fig. 2D**, **Supplementary Fig. 1**). The average reduction for the responsive HLA-A^∗^02:01 donors was 25-fold from α to α29 (n=7, p<0.03) and 30-fold from β to p29 (n=8; p<0.03; **Supplementary Fig. 1**). The predicted binding affinity to MHC class II was also decreased for α2 and β2 epitopes, although the experimental significance of this alteration is unknown.

We then generated modified Cas9 constructs by mutating the second residue of peptide α (L241G; Cas9-α2), peptide β (L616G; Cas9-β2), or both (Cas9-α2β2). To measure the effect of mutating the anchor residue of the immunogenic epitopes on T cell recognition of the Cas9 protein, we transiently transfected healthy donor B cell APCs with mRNA encoding wild type Cas9 (WT-Cas9), Cas9-α2, Cas9-β2, or Cas9-α2β2. Protein expression was confirmed by Western blot and the levels were comparable for all four constructs (data not shown). The T cell response measured by IFN-γ ELISpot after coculturing of transfected APCs with autologous PBMCs was significantly decreased for the modified Cas9 proteins (**Fig. 2E**). Introduction of the β2 mutation was the most effective in reducing T cell immunogenicity (5.5-fold, p<0.0001). This mutation in the REC1 domain (**Fig. 1E and 3A**) is not located in any of the two regions that are absolutely essential for DNA cleavage, the repeat-interacting (97–150) and the anti-repeat-interacting (312–409) regions ^31^. These results demonstrate that mutating the anchor amino acid of a highly immunogenic epitope can influence the overall immunogenicity of Cas9. Thus, engineering Cas9 variants with reduced immunogenicity potential can be used in conjunction with other strategies for safer CRISPR therapies and even possibly reduce the dosage of systemic immunosuppression needed for patients.

**Figure 3.**
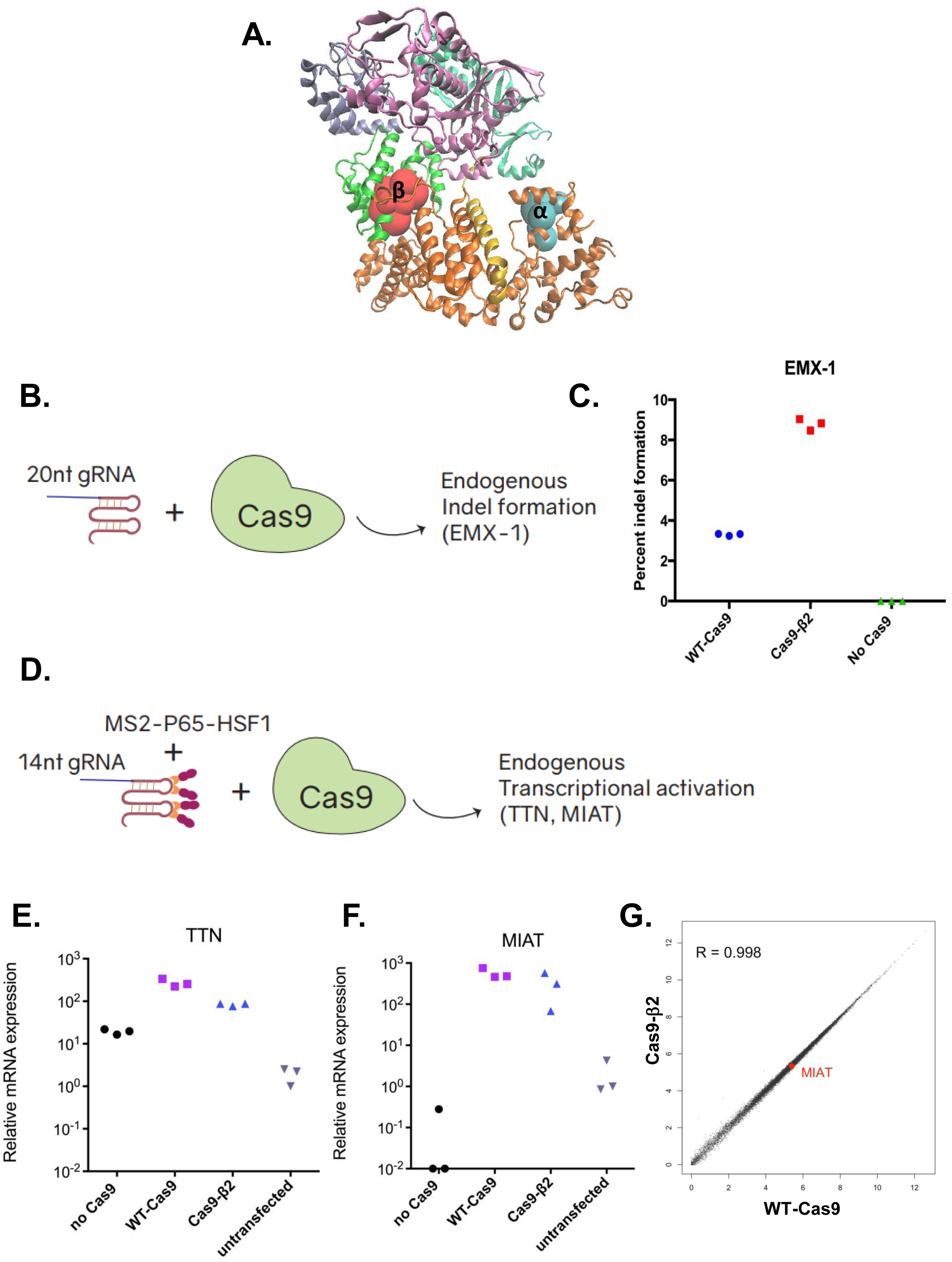
Mutated SpCas9 protein (Cas9-β2) retains its function and specificity. **A.** 3D structure of the SpCas9 protein, showing the location of the identified immunodominant epitopes α and β. **B.** Schematic of the experiment assessing mutagenesis capacity of Cas9-β2. Cells were transfected with either WT-Cas9, Cas9-β2, or an empty plasmid as well as 20nt gRNA targeting *EMX*-*1* locus. 72 hrs after transfection, percent cleavage was assessed by DNA extraction and illumina sequencing. **C.** Percentage of indel formation in *EMX*-*1* locus. Data represent mean +/- SD of three individual transfections. **D.** Schematic of the experiment assessing gRNA binding, DNA targeting and transcriptional modulation with Cas9-β2. Cells were transfected with either WT-Cas9, Cas9-β2, or an empty plasmid as well as 14nt gRNA targeting *TTN* or *MIAT* in the presence of MS2-P65-HSF1 (transcriptional modulation). 72 hrs after transfection, mRNA was assessed by qRT-PCR. **E,F.** Shown is the mRNA level relative to an untransfected control experiment. Each individual dot represents an individual transfection. **G.** Mean expression levels of 24,078 protein-coding and non-coding RNA genes for WT-Cas9 and Cas9-β2 (each in duplicate) are shown. For visualization purposes, the values were transformed to a log_2_(CPM+1) scale. *MIAT,* the gRNA target gene, is highlighted in red, and R denotes Pearson correlation coefficient between two groups.

We then tested the function of Cas9-β2 in comparison with WT-Cas9 in the context of DNA cleavage and transcriptional modulation. To examine the nuclease activity of Cas9-β2 and compare with WT-Cas9, we targeted Cas9-β2 or WT-Cas9 to an endogenous locus (*EMX*-*1*) and measured percent indel formation (**Fig. 3B, C**). Our data demonstrate that Cas9-β2 retains nuclease capacity in the locus we studied as well as on a synthetic promoter (**Fig. 3C**, **Supplementary Fig. 2A**).

Next, we determined whether Cas9-β2 can successfully recognize and bind its target DNA leading to transcriptional modulation. We first tested this in the context of enhanced transgene expression from a synthetic CRISPR responsive promoter in HEK293 cells using 14nt gRNAs and aptamer-mediated recruitment of transcriptional modulators similar to what we had shown before (**Supplementary Fig. 2B**). Having shown successful transgene activation, we then investigated whether this variant retains such capacity within the chromosomal contexts of endogenous genes. We transfected the cells with plasmids encoding Cas9-β2 or WT-Cas9 and 14nt gRNAs against two different endogenous genes (*TTN* and *MIAT*). qRT-PCR analysis showed that this variant successfully led to target gene expression (**Fig. 3D-F**). To further characterize Cas9-β2 specificity, we performed genome-wide RNA sequencing after targeting Cas9-β2 or WT-Cas9 to the *MIAT* locus for transcriptional activation. The results demonstrated no significant increase in undesired off-target activity by Cas9-β2 as compared to WT-Cas9 (**Fig. 3G**).

To show the extensibility of our approach, we tested the function of Cas9-α2, that has a mutation located in the REC2 domain (**Fig. 1E, 3A**). Cas9-α2 also demonstrated DNA cleavage and transcriptional modulation functionality comparable with WT-Cas9 (**Supplementary Fig. 3 A-E**). This is consistent with a previous study which showed that Cas9 with a deleted REC2 domain retains its nuclease activity ^31^. When T cells were stimulated with APCs spiked with peptide α2, the percentage of CD8+ CD137+ T cells (a marker of T cell activation ^33^) was decreased by 2.3-fold as compared to WT peptide α stimulation (**Supplementary Fig. 3F**).

## Discussion

The detection of pre-existing B cell and T cell immunity to the most widely used nuclease ortholog of the CRISPR/Cas9 tool in a significant proportion of healthy humans confirms previous studies in mice ^9,10^ and sheds light on the need for more studies of the immunological risks of this system. The CD8+ T cell immunity we observed is likely memory responses, as they are observed without ex vivo stimulation. However, following 18 days of T cell stimulation by peptides α or β, expansion of naïve T cells is not precluded. This suggests that the expression of Cas9 in naïve individuals may trigger a T cell response that could prevent subsequent administration. This could be avoided by switching to Cas9 orthologs from other bacterial species, but attention needs to be given to individual and distinct immune repertoires. This can be difficult given the epitope conservation across Cas9 proteins from multiple *Streptococcus* species and resemblance to sequences from other bacterial proteins such as the common pathogen *N. meningitidis*, that asymptomatically colonizes the nasopharynx in 10% of the population ^34^. Therefore, selective deimmunization (immunosilencing) of Cas9 can represent an attractive alternative, particularly in patients where high-dose systemic immunosuppression is contraindicated, such as in patients with chronic infectious diseases.

This strategy can be important in particular when longer term expression of Cas9 will be desired. Conventional methods of deimmunizing non-human therapeutic proteins rely on trial-and-error mutagenesis or machine learning and often includes deletion of whole regions of the protein ^35-39^. Here, as a general principle, we show that alteration of one of the anchor residues of an immunodominant epitope abolished specific T cell recognition. However, HLA allotype diversity and the existence of numerous epitopes in the large Cas9 protein complicate the process of complete deimmunization. The overall impact of removal of select immunodominant epitopes remains to be seen; both reduction ^40^ and enhancement ^41^ of the immunogenicity of subdominant epitopes have been reported with similar approaches for other proteins.

The top binding T cell epitopes within Cas9 that are most promiscuous for common HLA class I and class II alleles have been recently predicted *in silico* using IEDB ^42^. However, this is the first study that experimentally validates predicted immunodominant epitopes. None of the epitopes we report overlap with the peptides previously predicted ^42^. This is not unsurprising since we restricted our analysis to one HLA haplotype. Additionally, improved algorithms are needed to predict epitopes that hold up in experimental validation, as we show here. The use of CRISPR/Cas9 in humans may eventually necessitate creating HLA type-specific Cas9 variants, particularly for applications that require long-term Cas9 expression.

Non-specific localized immune suppressive approaches, such as those used by tumor cells and some viruses may complement these strategies for complete deimmunization. One attractive strategy is the transient and inducible co-expression of programmed death-ligand 1 (PD-L1) or indoleamine 2,3-dioxygenase 1 (IDO1) activating gRNAs inside cells that express Cas9 to protect them against attack by T cells. Alternatively, antigen presentation can be blocked by viral proteins interfering with antigen presentation (VIPRs), such as the adenoviral E319K or US2 and US11 from the human cytomegalovirus ^43^ or molecules that inhibit proteasomal antigen processing such as the Epstein-Barr virus Gly-Ala repeat ^44^. Deimmunized Cas9 may be useful in reduction of the dosage of other immunomodulatory measures needed to be coadministered in patients, thus facilitating therapeutic CRISPR applications as we develop better understanding of the immunological consequences of this system.

## Acknowledgements

The authors thank Dr. Erich Sturgis for providing the serum samples and Dr. Diego Chowell for support with the computational prediction model. We thank all the other members of the Anderson, Kiani, and Ebrahimkhani labs for their assistance and insightful discussions. This work was supported by ASU institutional funds and the startup fund by the School of Biological and Health Systems Engineering of ASU. We thank the Center for Computational and Integrative Biology (CCIB) DNA core facility at Massachusetts General Hospital and Genomics and Bioinformatics (TCGB) core at UCLA for the DNA and RNA sequencing services.

## Author Contributions

S.R.F. and R.E. designed experiments, performed experiments, and analyzed data. F.M. generated Cas9 variant constructs, performed Cas9 functional analysis experiments, and analyzed data. S. Krishna generated predicted Cas9 epitopes and designed and assisted with the T cell experiments. J.G.P. analyzed RNA seq data. M.R.E. helped with the design of experiments and interpretation of data. S.K. and K.S.A. supervised this study. S.R.F., R.E., S.K., and K.S.A. wrote the manuscript with the support of all the other authors.

## Methods

### Detection of Cas9-Specific Serum Antibodies in Healthy Controls

Healthy control sera (n = 183) used in this study, and previously described ^45^, are a subset of a molecular epidemiology study of head and neck cancer at the MD Anderson Cancer Center, collected between January 2006 and September 2008. Samples were collected using a standardized sample collection protocol and stored at −80°C until use. Written informed consent was obtained from all participants under institutional review board approval. *S. pyogenes* lysate was prepared by sonication of bacterial pellets from overnight cultures of *S. pyogenes* ATCC 19615 in the presence of 1 pill of complete Protease Inhibitor (Sigma-Aldrich) after 3 cycles of freezing and thawing. Serum antibody detection was performed using ELISA. 96-well plates were coated with 20 μg/mL of recombinant *S. pyogenes* Cas9 nuclease (New England Biolabs, Ipswich, MA) or *S. pyogenes* lysate. Sera were diluted 1:50 in 10% *E. coli* lysate prepared in 5% milk-PBST (0.2% tween) ^46^, incubated with shaking for 2 hrs at room temperature, and added to the specified wells in duplicate. Horseradish peroxidase (HRP) anti-human IgG Abs (Jackson ImmunoResearch Laboratories, West Grove, PA) were added at 1:10,000, and detected using Supersignal ELISA Femto Chemiluminescent substrate (Thermo Fisher Scientific, Waltham, MA). Luminescence was detected as relative light units (RLU) on a Glomax 96 Microplate Luminometer (Promega, Madison, WI) at 425 nm. To establish cut-off values, a RLU ratio > (the mean + 2 standard deviations) of all samples with signal below the mean RLU (horizontal black lines) was designated positive (**Fig. 1A**, dotted and dashed lines for bacterial lysate and Cas9 protein, respectively).

### Cas9 candidate T cell epitope prediction

We used our previously described prediction strategies ^30,47^ to predict candidate Cas9 T cell epitopes. Briefly, we predicted MHC class I restricted 9-mer and 10-mer candidate epitopes derived from the Cas9 protein (Uniprot - Q99ZW2) for HLA A^∗^02:01. The protein reference sequence was entered into 5 different prediction algorithms; 3 MHC-binding: IEDB-consensus binding ^48^, NetMHCpan binding ^49^, Syfpeithi ^50^ and 2 antigen-processing algorithms: IEDB-consensus processing, ANN processing ^51^. The individual scores from each of the prediction algorithms were then normalized within the pool of predicted peptides after exclusion of poor binders as previously detailed ^30,47^, and the average normalized binding scores were used to rerank the candidate peptides. The top 38 candidate peptides (**Supplementary Table 1**) were selected for experimental testing. The IEDB consensus MHC-binding prediction algorithm (http://www.iedb.org/) was applied to obtain a list of high binding Cas9 peptides, each of which was assigned a normalized binding score (S_b_). The immunogenicity score (S_i_) was calculated for each peptide based on its amino acid hydrophobicity (ANN-Hydro) ^30^.

### Ex vivo stimulation and epitope mapping of Cas9 by ELISpot

All peripheral blood mononuclear cells (PBMCs) were obtained from healthy individuals with written informed consent under ASU’s Institutional Review Board. PBMCs were isolated from fresh heparinized blood by Ficoll–Hypaque (GE Healthcare, UK) density gradient centrifugation and stimulated as previously described ^47^. Briefly, predicted Cas9 peptides with S_b_ < 0.148 (N=38) were synthesized (> 80% purity) by Proimmune, UK. Each peptide was reconstituted at 1mg/mL in sterile PBS and pools were created by mixing 3-4 candidate peptides. Sterile multiscreen ELISpot plates (Merck Millipore, Billerica, MA, USA) were coated overnight with 5μg/well of anti-IFN-γ capture antibody (clone D1K, Mabtech, USA) diluted in sterile PBS. Frozen PBMCs were thawed rapidly and recombinant human IL-2 (20U/mL, R&D Systems) was added. They were then stimulated in triplicates with 10μg/mL Cas9 peptide pools (or individual peptides), pre-mixed CEF pool as a positive control (ProImmune, UK), or DMSO as a negative control in the anti-IFN-γ-coated ELISpot plates, (Merck Millipore, Billerica, MA, USA) and incubated in a 37°C, 5% CO_2_ incubator for 48 hrs. Plates were washed three times for 5 min each with ELISpot buffer (PBS + 0.5% FBS) and incubated with 1μg/mL anti-IFN-γ secondary detection antibody (clone 7-B6-1, Mabtech, USA) for 2 hrs at room temperature, washed and incubated with 1μg/mL Streptavidin ALP conjugate for 1 hr at room temperature. The wells were washed again with ELISpot buffer and spots were developed by incubating for 8-10 min with detection buffer (33μL NBT, 16.5μL BCIP, in 100mM Tris-HCl pH 9, 1mM MgCl_2_, 150mM NaCl). Plates were left to dry for 2 days and spots were read using the AID ELISpot reader (Autoimmun Diagnostika GmbH, Germany). The average number of spot forming units for each triplicate was calculated for each test peptide or peptide pool and subtracted from the background signal.

### Autologous APC generation from healthy individual PBMCs

Autologous CD40L-activated B cell APCs were generated from healthy donors by incubating whole PBMCs with irradiated (32 Gy) K562-cell line expressing human CD40L (KCD40L) at a ratio of 4:1 (800,000 PBMCs to 200,000 irradiated KCD40Ls) in each well. The cells were maintained in B cell media (BCM) consisting of IMDM (Gibco, USA), 10% heat-inactivated human serum (Gemini Bio Products, CA, USA), and Antibiotic-Antimycotic (Anti-Anti, Gibco, USA). BCM was supplemented with 10ng/mL recombinant human IL-4 (R&D Systems, MN, USA), 2μg/mL Cyclosporin A (Sigma-Aldrich, CA, USA), and insulin transferrin supplement (ITES, Lonza, MD, USA). APCs were re-stimulated with fresh irradiated KCD40Ls on days 5 and 10, after washing with PBS and expanding into a whole 24-well plate. After two weeks, APC purity was assessed by CD19+ CD86+ expressing cells using flow cytometry, and were used for T cell stimulation after >90% purity. APCs were either restimulated up to 4 weeks or cryopreserved for re-expansion as necessary.

### T cell stimulation by autologous APCs

Antigen-specific T cells were detected by stimulating healthy donor B cell APCs by either peptide pulsing of specific Cas9 epitopes, or by transfecting with mRNA encoding the whole WT or modified Cas9 proteins. Peptide pulsing of APCs was done under BCM 5% human serum, with recombinant IL-4. Transfection of APCs was done with primary P3 buffer in a Lonza 4D Nucleofector and program EO117 (Lonza, MD, USA) and incubated in BCM-10% human serum and IL-4. Twenty-four hrs later, on day 1, APCs were washed and incubated with thawed whole PBMCs at a ratio of 1:2 (200,000 APCs : 400,000 PBMCs) in a 24-well plate in BCM supplemented with 20U/mL recombinant human IL-2 (R&D Systems, MN, USA) and 5ng/mL IL-7 (R&D Systems, MN, USA). On day 5, partial media exchange was performed by replacing half the well with fresh BCM and IL-2. On day 10, fresh APCs were peptide pulsed in a new 24-well plate. On day 11, expanded T cells were restimulated with peptide-pulsed APCs similar to day 1. T cells were used for T cell assays or immunophenotyped after day 18.

### Flow cytometry staining for T cells

Cells were washed once in MACS buffer (containing PBS, 1% BSA, 0.5mM EDTA), centrifuged at 550g for 5 min and re-suspended in 200μL MACS buffer. Cells were stained in 100μL of staining buffer containing anti-CD137, conjugated with phycoerythrin (PE, clone 4B4-1; BD Biosciences, USA), anti-CD8-PC5 (clone B9.11; Beckman Coulter 1:100), anti-CD4 (clone SK3; BioLegend, 1:200), anti-CD14 (clone 63D3; BioLegend, 1:200), and anti-CD19 (clone HIB19; BioLegend,1:200), all conjugated to Fluorescein isothiocyanate (FITC) for exclusion gates, for 30 min on ice. Samples were covered and incubated for 30 min on ice, washed twice in PBS, and resuspended in 1mL PBS prior to analysis.

### Pentamer staining for T cell immunophenotyping

The following HLA-A^∗^02:01 PE-conjugated Cas9 pentamers were obtained from ProImmune: F2A-D-CUS-A^∗^02:01-ILEDIVLTL-Pentamer, 007-Influenza A MP 58-66-GILGFVFTL-Pentamer. T cells were washed twice in MACS buffer with 5% human serum and centrifuged at 550g for 5 min each time. They were then re-suspended in 100μL staining buffer (MACS buffer, with 5% human serum and 1mM Dasatanib (ThermoFisher Scientific, MA, USA). Each of the pentamers was added to resuspended T cells, stimulated with the respective peptide or APCs at a concentration of 1:100. Samples were incubated at room temperature for 30 min in the dark, then washed twice in MACS buffer. Cells were stained in 100μL MACS buffer with anti-CD8-PC5, anti-CD4-FITC, anti-CD14-FITC, and anti-CD19-FITC for exclusion gates, Samples were then washed twice with PBS and analyzed by flow cytometry. For flow cytometric analysis, all samples were acquired with Attune flow cytometer (ThermoFisher Scientific, MA, USA) and analyzed using the Attune software. Gates for expression of different markers and pentamers were determined based on flow minus one (FMO) samples for each color after doublet discrimination (**Supplementary Fig. 4**). Percentages from each of the gated populations were used for the analysis.

### Vector Design and Construction

#### Modified Cas9 plasmids

- Human codon-optimized *Streptococcus pyogenes* Cas9 sequence was amplified from pSpCas9 (pX330; Addgene plasmid ID: 42230), using forward and reverse primers and inserted within gateway entry vectors using golden gate reaction. Desired mutations were designed within gBlocks (Integrated DNA Technologies). The gblocks and amplicons were then cloned into entry vectors using golden gate reaction. All the primers and gblocks sequences are listed in supplementary notes. Next, the Cas9 vectors and CAG promoter cassettes were cloned into an appropriate gateway destination vector via LR reaction (Invitrogen).

#### U6-sgRNA-MS2 plasmids

- These plasmids were constructed by inserting either 14bp or 20bp spacers of gRNAs (supplementary note) into sgRNA (MS2) cloning backbone (Addgene plasmid ID: 61424) at BbsI site. All the gRNA sequences are listed in supplementary note.

### Cell culture for endogenous target mutation and activation

HEK293FT cell line was purchased from ATCC and maintained in Dulbecco’s modified Eagle’s medium (DMEM - Life Technologies) containing 10% fetal bovine serum (FBS - Life Technologies), 2mM glutamine, 1mM sodium pyruvate (Life Technologies) and 1% penicillin-streptomycin (Life Technologies) in incubators at 37°C and 5% CO_2_. Polyethylenimine (PEI) was used to transfect HEK293FT cells seeded into 24-well plates. Transfection complexes were prepared according to manufacturer’s instructions.

### Fluorescent Reporter Assay for Quantifying Cas9 Function

HEK293FT cells were co-transfected with 10ng gRNA, 200ng Cas9 constructs, 100ng reporter plasmid and 25ng EBFP2 expressing plasmid as the transfection control. Fluorescent reporter experiments were performed 48 hrs after transfection. Flow cytometry data was analyzed using FlowJo. Cells were gated for positive EBFP expression to remove the un-transfected cells from the analysis (**Supplementary Fig. 4**). Untransfected controls were included in each experiment.

### Quantitative RT-PCR Analysis

HEK293FT cells were co-transfected with 10ng gRNA, 200ng Cas9 constructs, 100ng MS2-P65-HSF1 (Addgene plasmid ID: 61423) and 25ng transfection control. Cells were lysed, and RNA was extracted using RNeasy Plus mini kit (Qiagen) 72 hrs post transfection, followed by cDNA synthesis using the High-Capacity RNA-to-cDNA Kit (Thermo fisher). qRT-PCR was performed using SYBR Green PCR Master Mix (Thermo fisher). All analyses were normalized to 18S rRNA (ΔCt) and fold changes were calculated against un-transfected controls (2^−ΔΔCt^). Primer sequences for qPCR are listed in supplementary notes.

### Endogenous Indel Analysis

HEK293FT cells were co-transfected with 200ng of Cas9 plasmids, 10ng of gRNA coding cassette and and 25ng transfection control. 72 hrs later, transfected cells were dissociated and spun down at 200 g for 5 min at room temperature. Genomic DNA was extracted using 50 μl of QuickExtract DNA extraction solution (Epicentre) according to the manufacturer’s instructions. Genomic DNA was amplified by PCR using primers flanking the targeted region. Illumina Tru-Seq library was created by ligating partial adaptors and a unique barcode to the DNA samples. Next, a small number of PCR cycles was performed to complete the partial adaptors. Equal amounts of each sample were then pooled and sequenced on Illumina Tru-Seq platform with 2×150 run parameters, which yielded approximately 80,000 reads per sample. Sequencing was performed using a 2×150 paired-end (PE) configuration by CCIB DNA Core Facility at Massachusetts General Hospital (Cambridge, MA, USA). The reads were aligned to the target gene reference in *Mus musculus* genome using Geneious software, 9-1-5. To detect the indels (insertions and deletions of nucleic acid sequence at the site of double-strand break), each mutation was evaluated carefully in order to exclude the ones that are caused by sequencing error or any off-target mutation. The variant frequencies (percentage to total) assigned to each read containing indels were summed up. i.e. indel percentage = total number of indel containing reads/ total number of reads. The minimum number of analyzed reads per sample was 70,000.

### RNA Sequencing for Quantifying Activator Specificity

HEK293FT cells were co-transfected with 10ng gRNA for MIAT locus, 200ng Cas9 constructs, 100ng MS2-P65-HSF1 (Addgene plasmid ID: 61423) and 25ng transfection control. Total RNA was extracted 72 hrs post transfection using RNeasy Plus mini kit (Qiagen) and sent to UCLA TCGB core on dry ice. Ribosomal RNA depletion, and single read library preparation were performed at UCLA core followed by RNA sequencing using NextSeq500. Coverage was 14 million reads per sample. FASTQ files with single-ended 75bp reads were then aligned to the human GRCh38 reference genome sequence (Ensembl release 90) with STAR ^52^, and uniquely-mapped read counts (an average of 14.8 million reads per sample) were obtained with Cufflink ^53^. The read counts for each sample were then normalized for the library size to CPM (counts per million reads) with edgeR ^54^. Custom R scripts were then used to generate plots.

---

**Supplementary Table 1.**
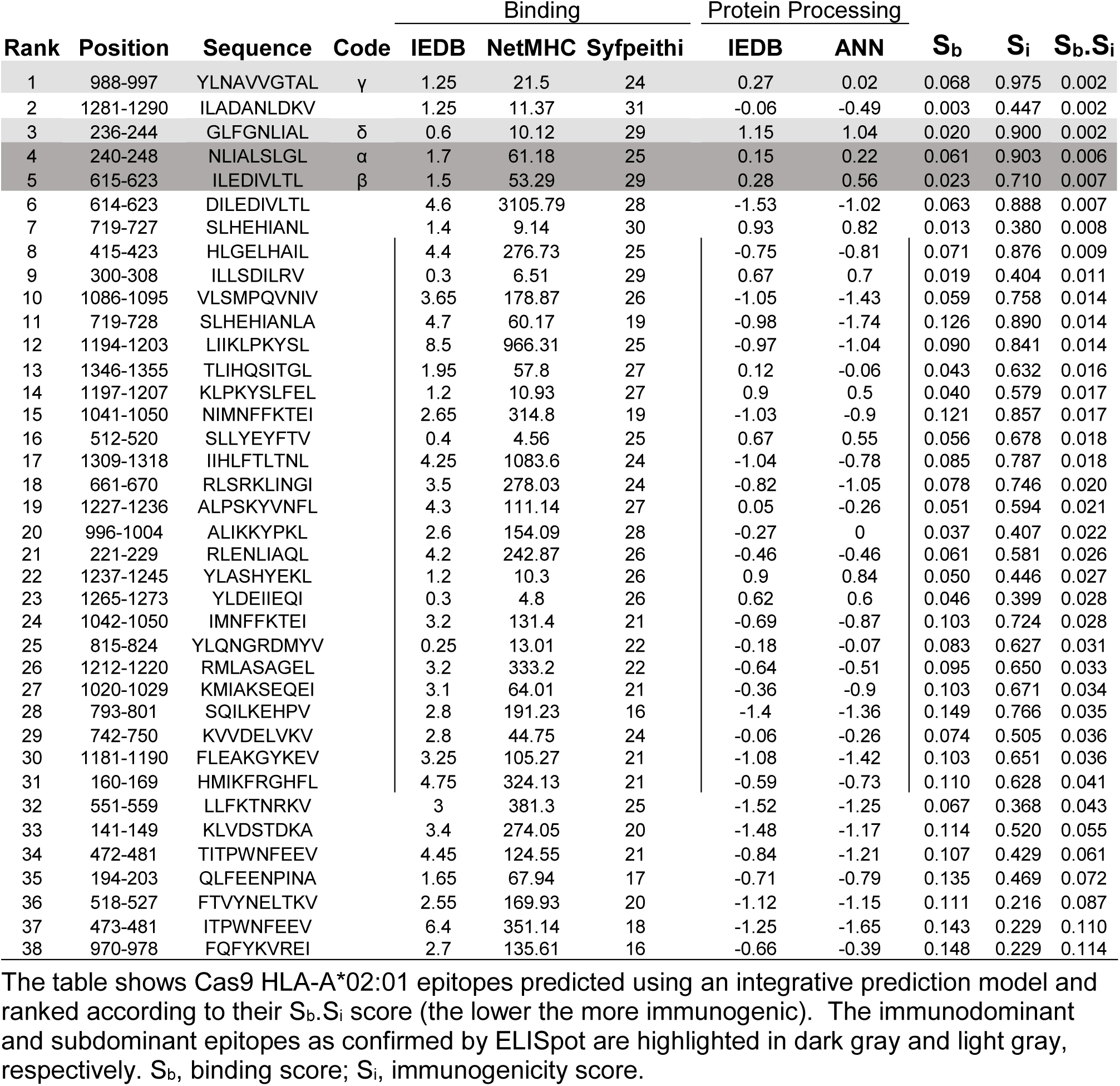
Predicted Cas9 immunogenic T cell epitopes

**Supplementary Table 3.**
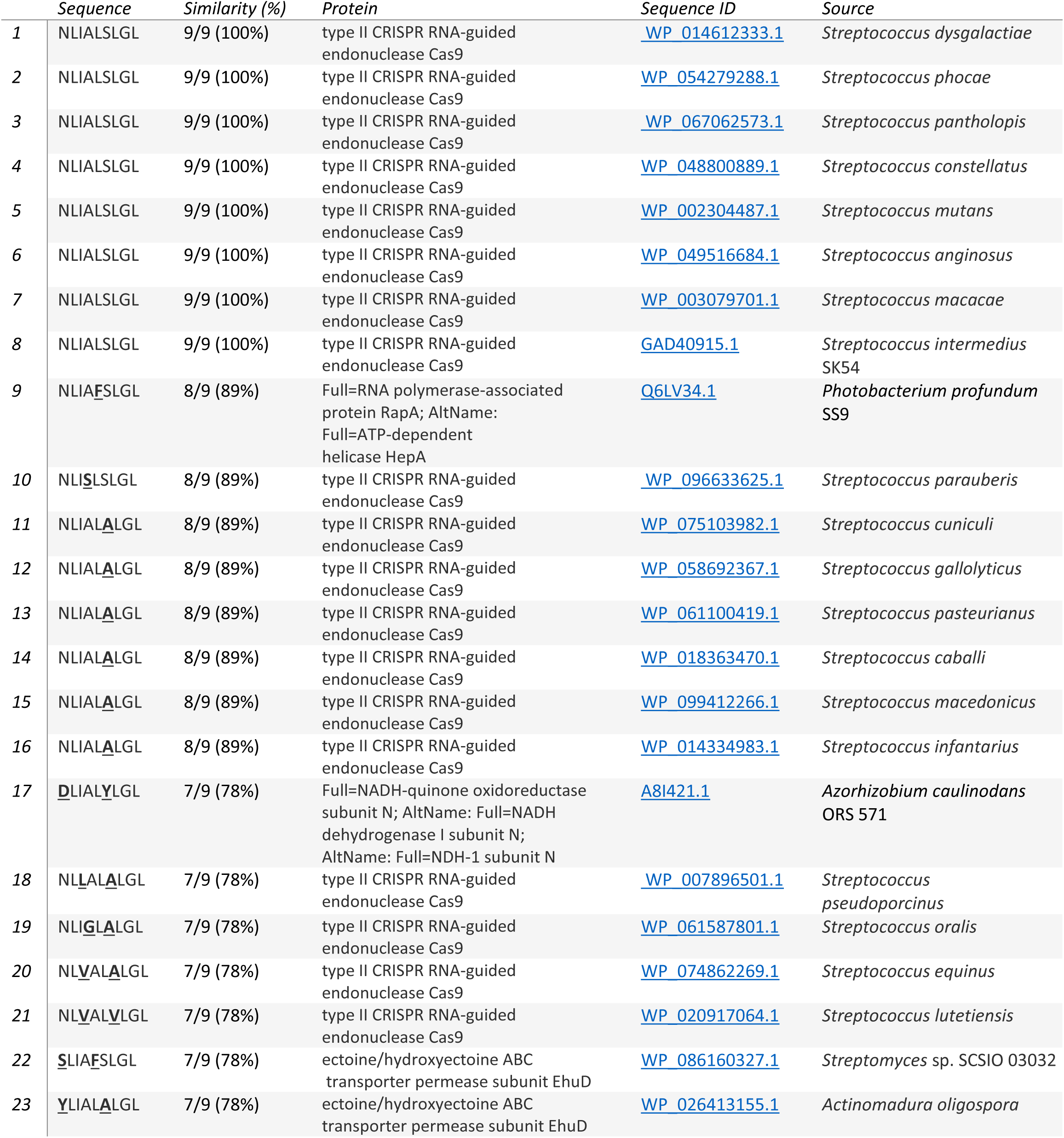
Sequence Homology of epitope α to amino acid sequences from known proteins.

**Supplementary Table 4.**
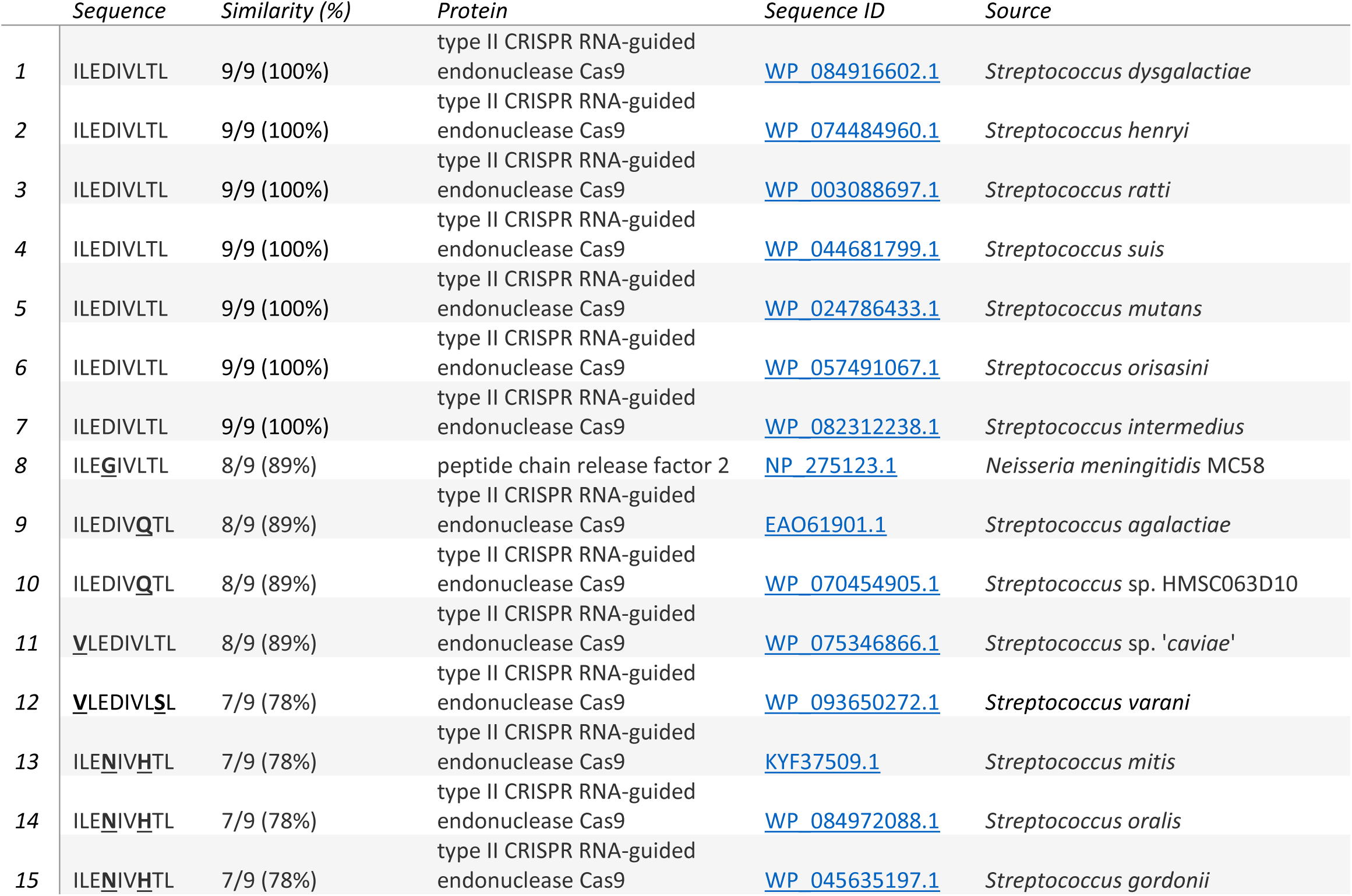
Sequence Homology of epitope β to amino acid sequences from known proteins.

|  | Sequence | Similarity (%) | Protein | Sequence ID | Source |
| --- | --- | --- | --- | --- | --- |
| 1 | ILEDIVLTL | 9/9 (100%) | type II CRISPR RNA-guided endonuclease Cas9 | WP_084916602.1 | *Streptococcus dysgalactiae* |
| 2 | ILEDIVLTL | 9/9 (100%) | type II CRISPR RNA-guided endonuclease Cas9 | WP_074484960.1 | *Streptococcus henryi* |
| 3 | ILEDIVLTL | 9/9 (100%) | type II CRISPR RNA-guided endonuclease Cas9 | WP_003088697.1 | *Streptococcus rattii* |
| 4 | ILEDIVLTL | 9/9 (100%) | type II CRISPR RNA-guided endonuclease Cas9 | WP_044681799.1 | *Streptococcus suis* |
| 5 | ILEDIVLTL | 9/9 (100%) | type II CRISPR RNA-guided endonuclease Cas9 | WP_024786433.1 | *Streptococcus mutans* |
| 6 | ILEDIVLTL | 9/9 (100%) | type II CRISPR RNA-guided endonuclease Cas9 | WP_057491067.1 | *Streptococcus orisasinii* |
| 7 | ILEDIVLTL | 9/9 (100%) | type II CRISPR RNA-guided endonuclease Cas9 | WP_082312238.1 | *Streptococcus intermedius* |
| 8 | I**L**E**G**IVLTL | 8/9 (89%) | peptide chain release factor 2 | NP_275123.1 | *Neisseria meningitidis* MC58 |
| 9 | ILEDIV**Q**TL | 8/9 (89%) | type II CRISPR RNA-guided endonuclease Cas9 | EAO61901.1 | *Streptococcus agalactiae* |
| 10 | ILEDIV**Q**TL | 8/9 (89%) | type II CRISPR RNA-guided endonuclease Cas9 | WP_070454905.1 | *Streptococcus* sp. HMSC063D10 |
| 11 | **V**LEDIVLTL | 8/9 (89%) | type II CRISPR RNA-guided endonuclease Cas9 | WP_075346866.1 | *Streptococcus* sp. 'caviae' |
| 12 | **V**LEDIV**L**SL | 7/9 (78%) | type II CRISPR RNA-guided endonuclease Cas9 | WP_093650272.1 | *Streptococcus varani* |
| 13 | I**L**E**N**IV**H**TL | 7/9 (78%) | type II CRISPR RNA-guided endonuclease Cas9 | KYF37509.1 | *Streptococcus mitis* |
| 14 | I**L**E**N**IV**H**TL | 7/9 (78%) | type II CRISPR RNA-guided endonuclease Cas9 | WP_084972088.1 | *Streptococcus oralis* |
| 15 | I**L**E**N**IV**H**TL | 7/9 (78%) | type II CRISPR RNA-guided endonuclease Cas9 | WP_045635197.1 | *Streptococcus gordonii* |

**Supplementary Figure 1.**
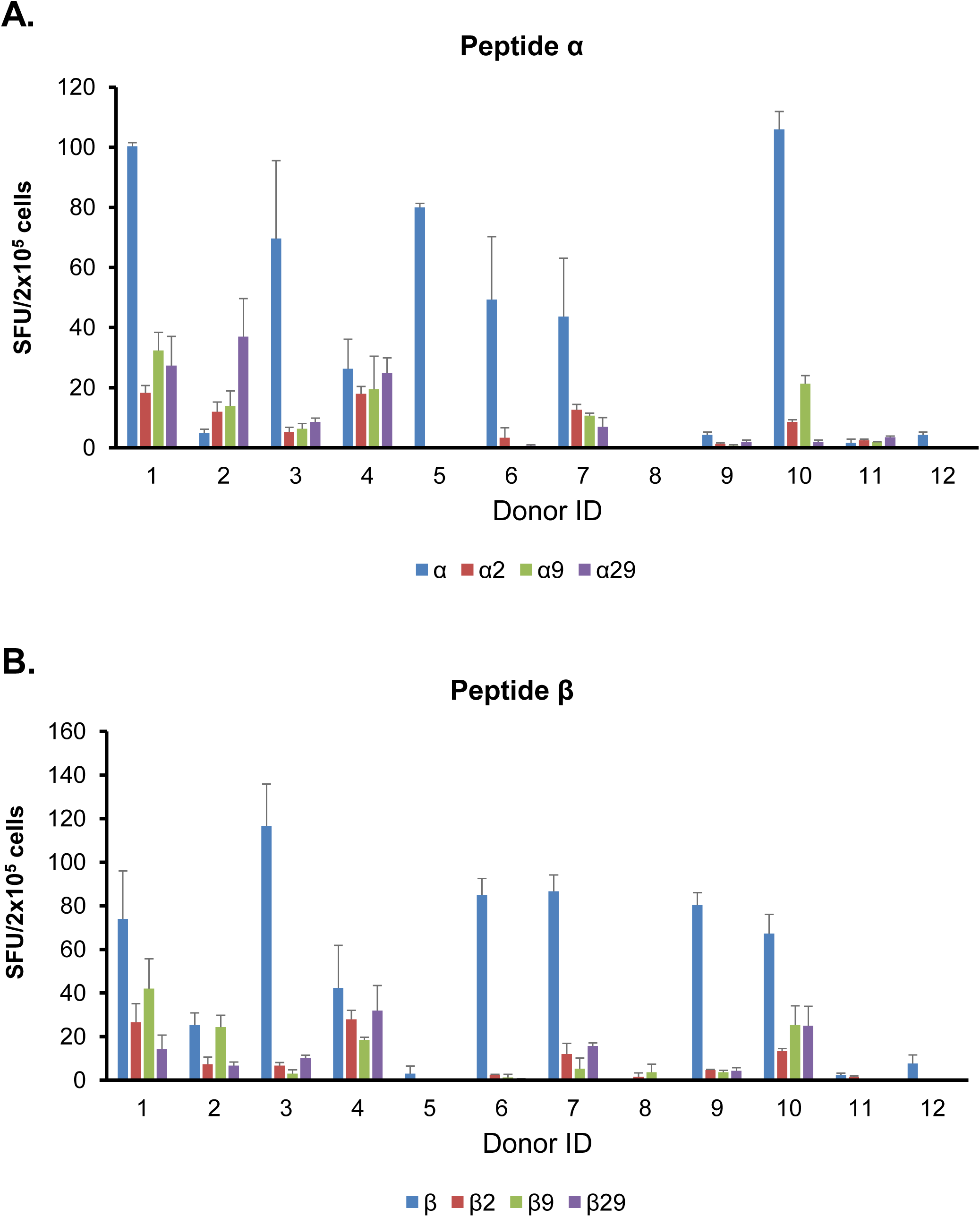
Reduced T cell response to epitopes α and β after mutation of the anchor residues. **A and B.** IFN-γ ELISpot for 12 healthy donor PBMCs stimulated with wild type or mutated peptide α (**A**) or β (**B**). Data represent mean +/- SEM.

**Supplementary Figure 2.**
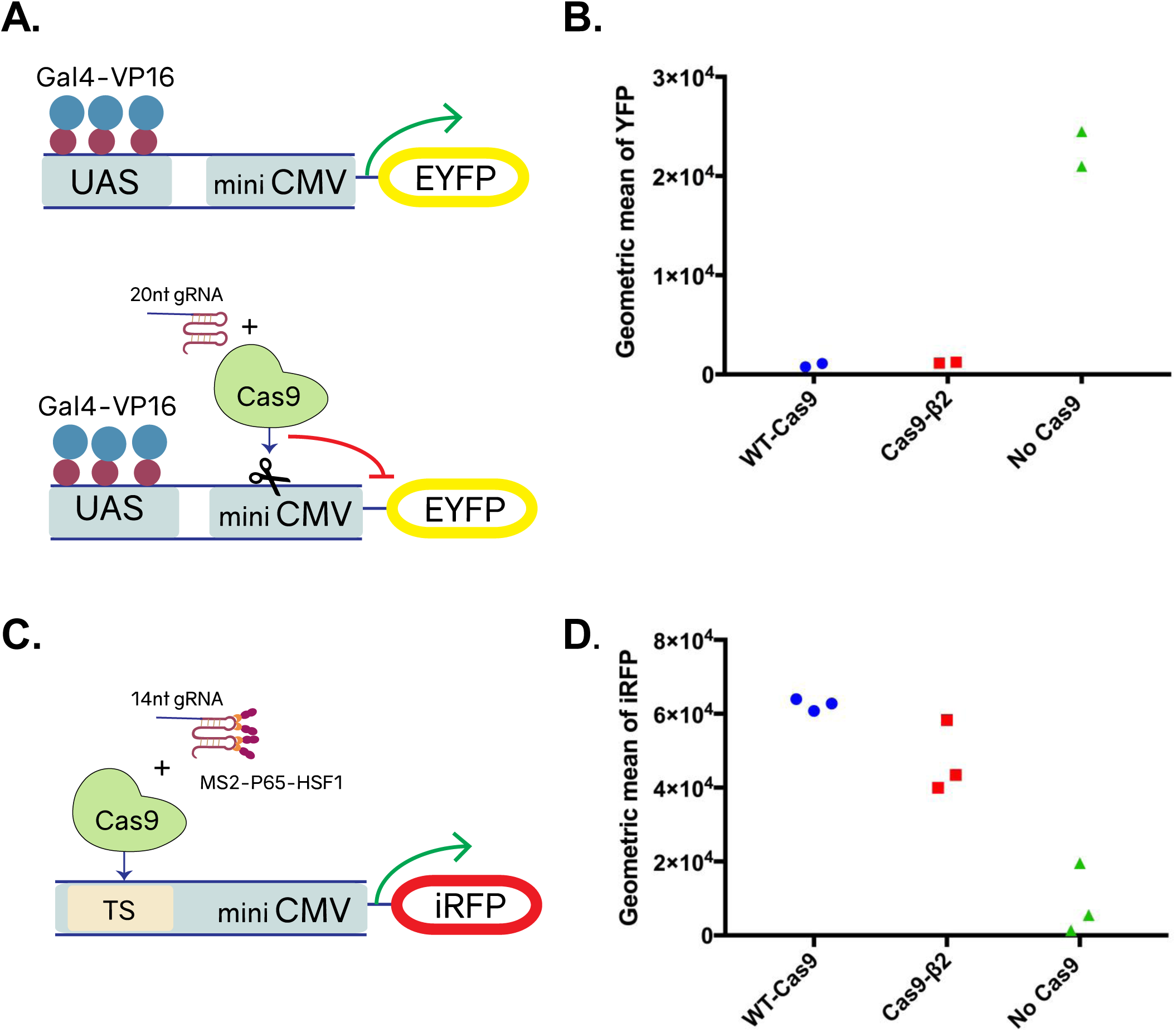
**A.** Schematic of the experiment assessing Cas9-β2 cleavage capacity at a synthetic promoter. Cells were transfected with either WT-Cas9, Cas9-β2 or an empty plasmid as well as 20nt gRNA targeting a synthetic CRISPR promoter that harbors two gRNA target sites flanking a mini-CMV promoter. The targeting and cleavage at the promoter should disrupt the promoter and decrease EYFP expression. **B.** Each individual dot represents EYFP expression 48 hrs after transfection in cells expressing >2×10^2^ A.U. of a transfection marker measured by flow cytometry (n=2 individual transfections represented by individual dots). **C.** Schematic of the experiment assessing Cas9-β2 transcriptional activation capacity at a synthetic promoter. Cells were transfected with either WT-Cas9, Cas9-β2 or an empty plasmid as well as aptamer binding transcriptional activation domains, and a 14nt gRNA targeting a synthetic CRISPR promoter that harbors multiple target sites upstream of a mini-CMV promoter. Targeting at the promoter should enable iRFP expression. **D.** Each individual dot shows iRFP expression 48 hrs after transfection in cells expressing >2×10^2^ A.U. of a transfection marker measured by flow cytometry (n=3 individual transfections represented by individual dots, n=2 for the no Cas9 group).

**Supplementary Figure 3.**
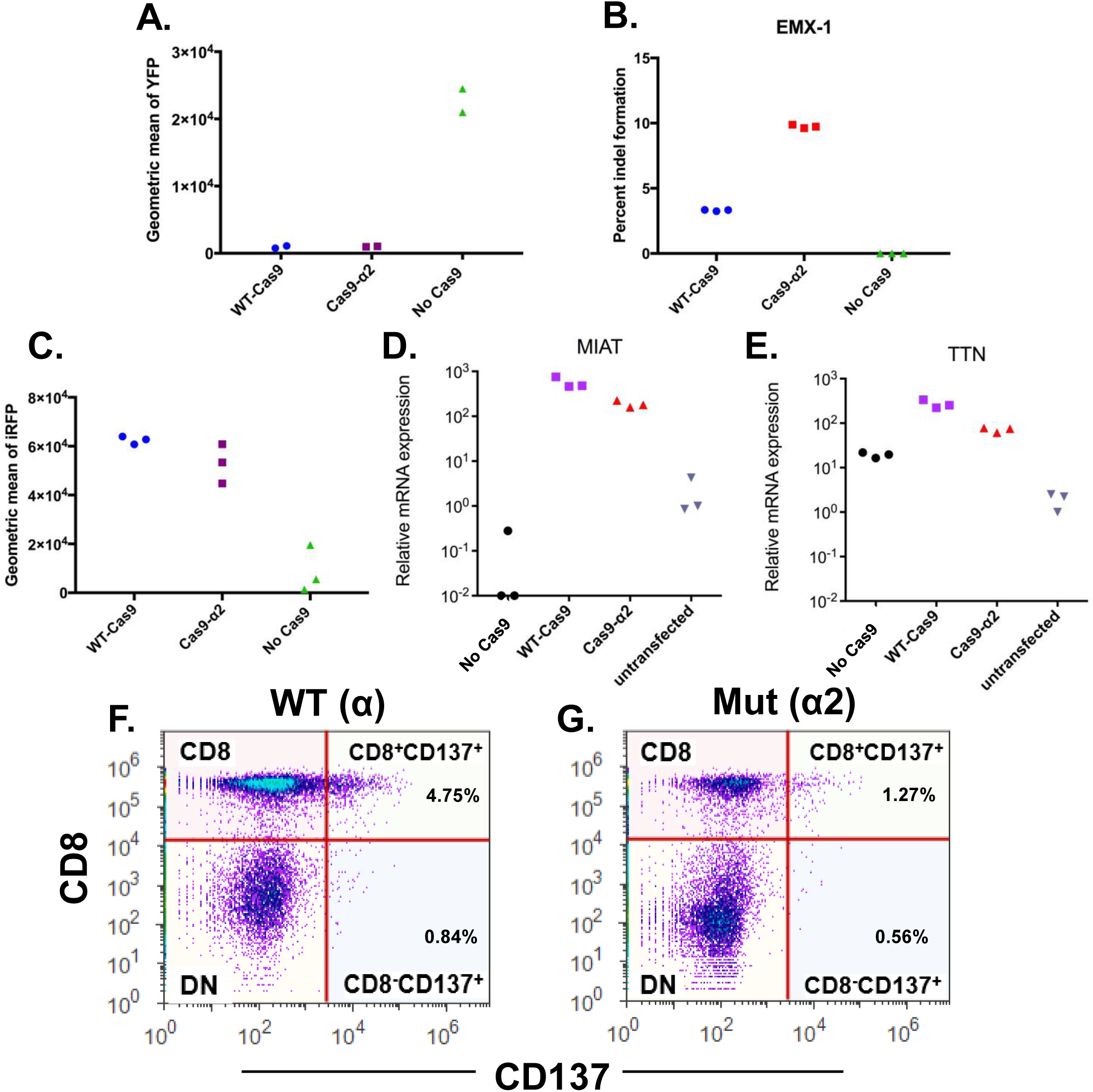
**A.** Analysis of the mutagenesis capacity of Cas9-α2 as compared to WT-Cas9 in a synthetic promoter. Each individual dot shows EYFP expression 48 hrs after transfection in cells expressing >2×10^2^ A.U. of a transfection marker measured by flow cytometry (n=2 individual transfections, represented by individual dots). **B.** Mutagenesis in the endogenous *EMX*-*1* locus. Percentage of indel formation in the *EMX-1 locus.* Dots represent three individual transfections **C.** Transcriptional modulation by Cas9-α2 at a synthetic promoter. Each individual dot shows iRFP expression 48 hrs after transfection in cells expressing >2×10^2^ A.U. of a transfection marker measured by flow cytometry (n=3 individual transfections, n=2 for no Cas9 group represented by dots). **D, E.** Shown is the mRNA level relative to an untransfected control experiment. Each individual dot represents an individual transfection. Note for A and C WT-Cas9 and no Cas9 data are also reported in Supplementary Fig.2. For B-E. WT-Cas9 and no Cas9 data are also reported in Fig.3. **F.** Activated CD8+CD137+ T cells detected in PBMCs stimulated with peptide α or peptide α2 (data representative of 3 individual stimulations).

**Supplementary Figure 4.**
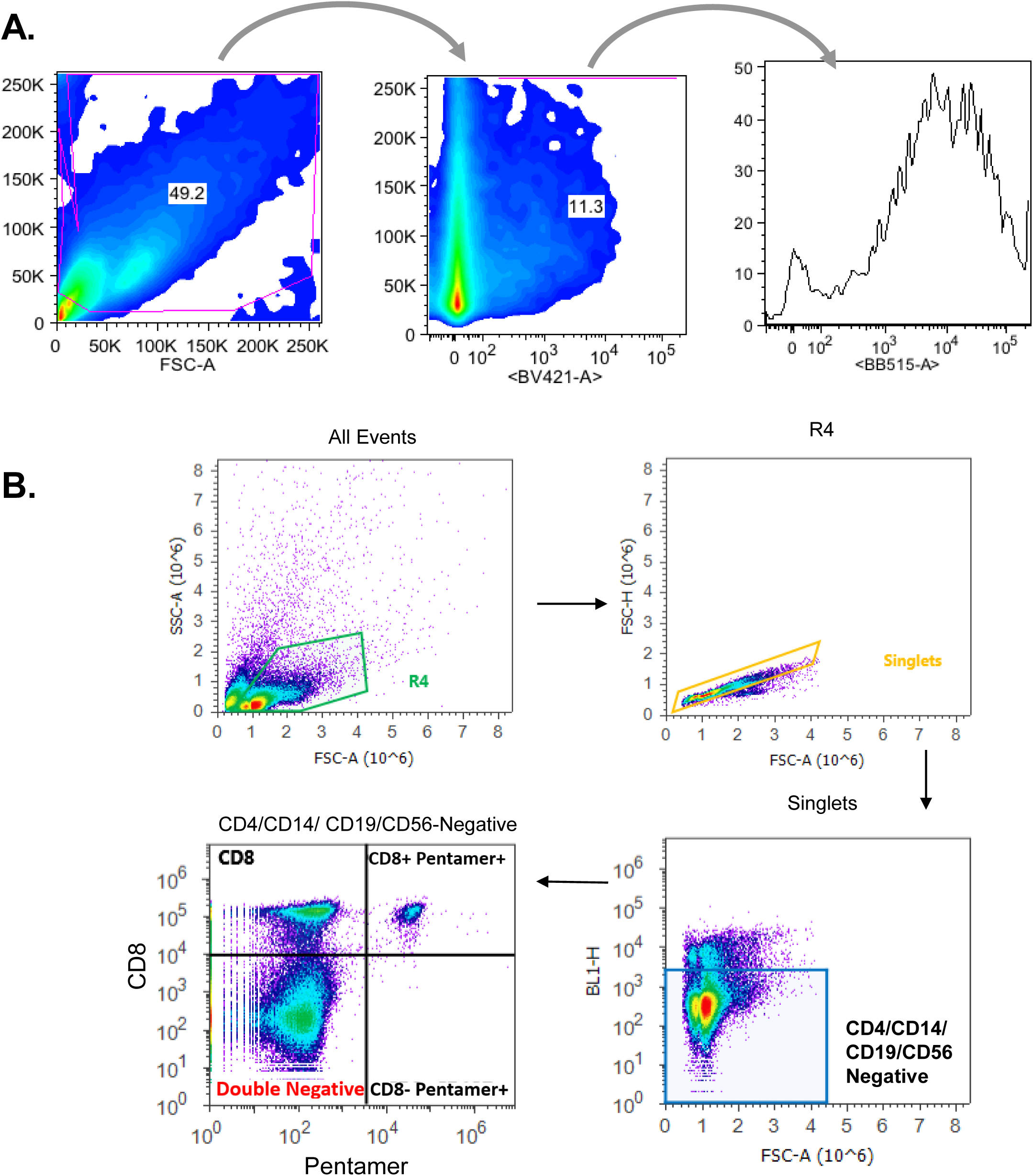
**A.** Representative flow cytometry gating for analysis of Cas9 function on synthetic promoters. Cells are gated based on Forward (FSC) and Side Scatter (SSCs). Transfected population then is gated based on expression of EBFP (BV421-A) more than 2×10^2^. The geometric mean of the output (EYFP) or iRFP is determined in this population. **B.** Flow cytometry gating for analysis of Cas9 pentamer+ CD8+ T lymphocytes. Cells are gated based on FSC and SSC and negatively gated on CD4/CD14/CD19/CD56. The CD8-, CD8+, and Cas9 pentamer+ population are shown (bottom right).

## Supplementary notes

### Sequences of Cas9

#### Cas9-β2

ATGGACTATAAGGACCACGACGGAGACTACAAGGATCATGATATTGATTACAAAGACGATGACGA TAAGATGGCCCCAAAGAAGAAGCGGAAGGTCGGTATCCACGGAGTCCCAGCAGCCGACAAGAAG TACAGCATCGGCCTGGACATCGGCACCAACTCTGTGGGCTGGGCCGTGATCACCGACGAGTACAA GGTGCCCAGCAAGAAATTCAAGGTGCTGGGCAACACCGACCGGCACAGCATCAAGAAGAACCTG ATCGGAGCCCTGCTGTTCGACAGCGGCGAAACAGCCGAGGCCACCCGGCTGAAGAGAACCGCCA GAAGAAGATACACCAGACGGAAGAACCGGATCTGCTATCTGCAAGAGATCTTCAGCAACGAGAT GGCCAAGGTGGACGACAGCTTCTTCCACAGACTGGAAGAGTCCTTCCTGGTGGAAGAGGATAAGA AGCACGAGCGGCACCCCATCTTCGGCAACATCGTGGACGAGGTGGCCTACCACGAGAAGTACCCC ACCATCTACCACCTGAGAAAGAAACTGGTGGACAGCACCGACAAGGCCGACCTGCGGCTGATCTA TCTGGCCCTGGCCCACATGATCAAGTTCCGGGGCCACTTCCTGATCGAGGGCGACCTGAACCCCG ACAACAGCGACGTGGACAAGCTGTTCATCCAGCTGGTGCAGACCTACAACCAGCTGTTCGAGGAA AACCCCATCAACGCCAGCGGCGTGGACGCCAAGGCCATCCTGTCTGCCAGACTGAGCAAGAGCAG ACGGCTGGAAAATCTGATCGCCCAGCTGCCCGGCGAGAAGAAGAATGGCCTGTTCGGAAACCTTA TTGCCCTGAGCCTGGGCCTGACCCCCAACTTCAAGAGCAACTTCGACCTGGCCGAGGATGCCAAA CTGCAGCTGAGCAAGGACACCTACGACGACGACCTGGACAACCTGCTGGCCCAGATCGGCGACCA GTACGCCGACCTGTTTCTGGCCGCCAAGAACCTGTCCGACGCCATCCTGCTGAGCGACATCCTGAG AGTGAACACCGAGATCACCAAGGCCCCCCTGAGCGCCTCTATGATCAAGAGATACGACGAGCACC ACCAGGACCTGACCCTGCTGAAAGCTCTCGTGCGGCAGCAGCTGCCTGAGAAGTACAAAGAGATT TTCTTCGACCAGAGCAAGAACGGCTACGCCGGCTACATTGACGGCGGAGCCAGCCAGGAAGAGTT CTACAAGTTCATCAAGCCCATCCTGGAAAAGATGGACGGCACCGAGGAACTGCTCGTGAAGCTGA ACAGAGAGGACCTGCTGCGGAAGCAGCGGACCTTCGACAACGGCAGCATCCCCCACCAGATCCAC CTGGGAGAGCTGCACGCCATTCTGCGGCGGCAGGAAGATTTTTACCCATTCCTGAAGGACAACCG GGAAAAGATCGAGAAGATCCTGACCTTCCGCATCCCCTACTACGTGGGCCCTCTGGCCAGGGGAA ACAGCAGATTCGCCTGGATGACCAGAAAGAGCGAGGAAACCATCACCCCCTGGAACTTCGAGGA AGTGGTGGACAAGGGCGCTTCCGCCCAGAGCTTCATCGAGCGGATGACCAACTTCGATAAGAACC TGCCCAACGAGAAGGTGCTGCCCAAGCACAGCCTGCTGTACGAGTACTTCACCGTGTATAACGAG CTGACCAAAGTGAAATACGTGACCGAGGGAATGAGAAAGCCCGCCTTCCTGAGCGGCGAGCAGA AAAAGGCCATCGTGGACCTGCTGTTCAAGACCAACCGGAAAGTGACCGTGAAGCAGCTGAAAGA GGACTACTTCAAGAAAATCGAGTGCTTCGACTCCGTGGAAATCTCCGGCGTGGAAGATCGGTTCA ACGCCTCCCTGGGCACATACCACGATCTGCTGAAAATTATCAAGGACAAGGACTTCCTGGACAAT GAGGAAAACGAGGACATTGGTGAAGATATCGTGCTGACCCTGACACTGTTTGAGGACAGAGAGAT GATCGAGGAACGGCTGAAAACCTATGCCCACCTGTTCGACGACAAAGTGATGAAGCAGCTGAAGC GGCGGAGATACACCGGCTGGGGCAGGCTGAGCCGGAAGCTGATCAACGGCATCCGGGACAAGCA GTCCGGCAAGACAATCCTGGATTTCCTGAAGTCCGACGGCTTCGCCAACAGAAACTTCATGCAGC TGATCCACGACGACAGCCTGACCTTTAAAGAGGACATCCAGAAAGCCCAGGTGTCCGGCCAGGGC GATAGCCTGCACGAGCACATTGCCAATCTGGCCGGCAGCCCCGCCATTAAGAAGGGCATCCTGCA GACAGTGAAGGTGGTGGACGAGCTCGTGAAAGTGATGGGCCGGCACAAGCCCGAGAACATCGTG ATCGAAATGGCCAGAGAGAACCAGACCACCCAGAAGGGACAGAAGAACAGCCGCGAGAGAATG AAGCGGATCGAAGAGGGCATCAAAGAGCTGGGCAGCCAGATCCTGAAAGAACACCCCGTGGAAA ACACCCAGCTGCAGAACGAGAAGCTGTACCTGTACTACCTGCAGAATGGGCGGGATATGTACGTG GACCAGGAACTGGACATCAACCGGCTGTCCGACTACGATGTGGACCATATCGTGCCTCAGAGCTT TCTGAAGGACGACTCCATCGACAACAAGGTGCTGACCAGAAGCGACAAGAACCGGGGCAAGAGC GACAACGTGCCCTCCGAAGAGGTCGTGAAGAAGATGAAGAACTACTGGCGGCAGCTGCTGAACG CCAAGCTGATTACCCAGAGAAAGTTCGACAATCTGACCAAGGCCGAGAGAGGCGGCCTGAGCGA ACTGGATAAGGCCGGCTTCATCAAGAGACAGCTGGTGGAAACCCGGCAGATCACAAAGCACGTG GCACAGATCCTGGACTCCCGGATGAACACTAAGTACGACGAGAATGACAAGCTGATCCGGGAAGT GAAAGTGATCACCCTGAAGTCCAAGCTGGTGTCCGATTTCCGGAAGGATTTCCAGTTTTACAAAGT GCGCGAGATCAACAACTACCACCACGCCCACGACGCCTACCTGAACGCCGTCGTGGGAACCGCCC TGATCAAAAAGTACCCTAAGCTGGAAAGCGAGTTCGTGTACGGCGACTACAAGGTGTACGACGTG CGGAAGATGATCGCCAAGAGCGAGCAGGAAATCGGCAAGGCTACCGCCAAGTACTTCTTCTACAG CAACATCATGAACTTTTTCAAGACCGAGATTACCCTGGCCAACGGCGAGATCCGGAAGCGGCCTC TGATCGAGACAAACGGCGAAACCGGGGAGATCGTGTGGGATAAGGGCCGGGATTTTGCCACCGTG CGGAAAGTGCTGAGCATGCCCCAAGTGAATATCGTGAAAAAGACCGAGGTGCAGACAGGCGGCT TCAGCAAAGAGTCTATCCTGCCCAAGAGGAACAGCGATAAGCTGATCGCCAGAAAGAAGGACTG GGACCCTAAGAAGTACGGCGGCTTCGACAGCCCCACCGTGGCCTATTCTGTGCTGGTGGTGGCCA AAGTGGAAAAGGGCAAGTCCAAGAAACTGAAGAGTGTGAAAGAGCTGCTGGGGATCACCATCAT GGAAAGAAGCAGCTTCGAGAAGAATCCCATCGACTTTCTGGAAGCCAAGGGCTACAAAGAAGTG AAAAAGGACCTGATCATCAAGCTGCCTAAGTACTCCCTGTTCGAGCTGGAAAACGGCCGGAAGAG AATGCTGGCCTCTGCCGGCGAACTGCAGAAGGGAAACGAACTGGCCCTGCCCTCCAAATATGTGA ACTTCCTGTACCTGGCCAGCCACTATGAGAAGCTGAAGGGCTCCCCCGAGGATAATGAGCAGAAA CAGCTGTTTGTGGAACAGCACAAGCACTACCTGGACGAGATCATCGAGCAGATCAGCGAGTTCTC CAAGAGAGTGATCCTGGCCGACGCTAATCTGGACAAAGTGCTGTCCGCCTACAACAAGCACCGGG ATAAGCCCATCAGAGAGCAGGCCGAGAATATCATCCACCTGTTTACCCTGACCAATCTGGGAGCC CCTGCCGCCTTCAAGTACTTTGACACCACCATCGACCGGAAGAGGTACACCAGCACCAAAGAGGT GCTGGACGCCACCCTGATCCACCAGAGCATCACCGGCCTGTACGAGACACGGATCGACCTGTCTC AGCTGGGAGGCGACAAAAGGCCGGCGGCCACGAAAAAGGCCGGCCAGGCAAAAAAGAAAAAGTAA

#### Cas9-α2

ATGGACTATAAGGACCACGACGGAGACTACAAGGATCATGATATTGATTACAAAGACGATGACGA TAAGATGGCCCCAAAGAAGAAGCGGAAGGTCGGTATCCACGGAGTCCCAGCAGCCGACAAGAAG TACAGCATCGGCCTGGACATCGGCACCAACTCTGTGGGCTGGGCCGTGATCACCGACGAGTACAA GGTGCCCAGCAAGAAATTCAAGGTGCTGGGCAACACCGACCGGCACAGCATCAAGAAGAACCTG ATCGGAGCCCTGCTGTTCGACAGCGGCGAAACAGCCGAGGCCACCCGGCTGAAGAGAACCGCCA GAAGAAGATACACCAGACGGAAGAACCGGATCTGCTATCTGCAAGAGATCTTCAGCAACGAGAT GGCCAAGGTGGACGACAGCTTCTTCCACAGACTGGAAGAGTCCTTCCTGGTGGAAGAGGATAAGA AGCACGAGCGGCACCCCATCTTCGGCAACATCGTGGACGAGGTGGCCTACCACGAGAAGTACCCC ACCATCTACCACCTGAGAAAGAAACTGGTGGACAGCACCGACAAGGCCGACCTGCGGCTGATCTA TCTGGCCCTGGCCCACATGATCAAGTTCCGGGGCCACTTCCTGATCGAGGGCGACCTGAACCCCG ACAACAGCGACGTGGACAAGCTGTTCATCCAGCTGGTGCAGACCTACAACCAGCTGTTCGAGGAA AACCCCATCAACGCCAGCGGCGTGGACGCCAAGGCCATCCTGTCTGCCAGACTGAGCAAGAGCAG ACGGCTGGAAAATCTGATCGCCCAGCTGCCCGGCGAGAAGAAGAATGGCCTGTTCGGAAACGGTA TTGCCCTGAGCCTGGGCCTGACCCCCAACTTCAAGAGCAACTTCGACCTGGCCGAGGATGCCAAA CTGCAGCTGAGCAAGGACACCTACGACGACGACCTGGACAACCTGCTGGCCCAGATCGGCGACCA GTACGCCGACCTGTTTCTGGCCGCCAAGAACCTGTCCGACGCCATCCTGCTGAGCGACATCCTGAG AGTGAACACCGAGATCACCAAGGCCCCCCTGAGCGCCTCTATGATCAAGAGATACGACGAGCACC ACCAGGACCTGACCCTGCTGAAAGCTCTCGTGCGGCAGCAGCTGCCTGAGAAGTACAAAGAGATT TTCTTCGACCAGAGCAAGAACGGCTACGCCGGCTACATTGACGGCGGAGCCAGCCAGGAAGAGTT CTACAAGTTCATCAAGCCCATCCTGGAAAAGATGGACGGCACCGAGGAACTGCTCGTGAAGCTGA ACAGAGAGGACCTGCTGCGGAAGCAGCGGACCTTCGACAACGGCAGCATCCCCCACCAGATCCAC CTGGGAGAGCTGCACGCCATTCTGCGGCGGCAGGAAGATTTTTACCCATTCCTGAAGGACAACCG GGAAAAGATCGAGAAGATCCTGACCTTCCGCATCCCCTACTACGTGGGCCCTCTGGCCAGGGGAA ACAGCAGATTCGCCTGGATGACCAGAAAGAGCGAGGAAACCATCACCCCCTGGAACTTCGAGGA AGTGGTGGACAAGGGCGCTTCCGCCCAGAGCTTCATCGAGCGGATGACCAACTTCGATAAGAACC TGCCCAACGAGAAGGTGCTGCCCAAGCACAGCCTGCTGTACGAGTACTTCACCGTGTATAACGAG CTGACCAAAGTGAAATACGTGACCGAGGGAATGAGAAAGCCCGCCTTCCTGAGCGGCGAGCAGA AAAAGGCCATCGTGGACCTGCTGTTCAAGACCAACCGGAAAGTGACCGTGAAGCAGCTGAAAGA GGACTACTTCAAGAAAATCGAGTGCTTCGACTCCGTGGAAATCTCCGGCGTGGAAGATCGGTTCA ACGCCTCCCTGGGCACATACCACGATCTGCTGAAAATTATCAAGGACAAGGACTTCCTGGACAAT GAGGAAAACGAGGACATTCTTGAAGATATCGTGCTGACCCTGACACTGTTTGAGGACAGAGAGAT GATCGAGGAACGGCTGAAAACCTATGCCCACCTGTTCGACGACAAAGTGATGAAGCAGCTGAAGC GGCGGAGATACACCGGCTGGGGCAGGCTGAGCCGGAAGCTGATCAACGGCATCCGGGACAAGCA GTCCGGCAAGACAATCCTGGATTTCCTGAAGTCCGACGGCTTCGCCAACAGAAACTTCATGCAGC TGATCCACGACGACAGCCTGACCTTTAAAGAGGACATCCAGAAAGCCCAGGTGTCCGGCCAGGGC GATAGCCTGCACGAGCACATTGCCAATCTGGCCGGCAGCCCCGCCATTAAGAAGGGCATCCTGCA GACAGTGAAGGTGGTGGACGAGCTCGTGAAAGTGATGGGCCGGCACAAGCCCGAGAACATCGTG ATCGAAATGGCCAGAGAGAACCAGACCACCCAGAAGGGACAGAAGAACAGCCGCGAGAGAATG AAGCGGATCGAAGAGGGCATCAAAGAGCTGGGCAGCCAGATCCTGAAAGAACACCCCGTGGAAA ACACCCAGCTGCAGAACGAGAAGCTGTACCTGTACTACCTGCAGAATGGGCGGGATATGTACGTG GACCAGGAACTGGACATCAACCGGCTGTCCGACTACGATGTGGACCATATCGTGCCTCAGAGCTT TCTGAAGGACGACTCCATCGACAACAAGGTGCTGACCAGAAGCGACAAGAACCGGGGCAAGAGC GACAACGTGCCCTCCGAAGAGGTCGTGAAGAAGATGAAGAACTACTGGCGGCAGCTGCTGAACG CCAAGCTGATTACCCAGAGAAAGTTCGACAATCTGACCAAGGCCGAGAGAGGCGGCCTGAGCGA ACTGGATAAGGCCGGCTTCATCAAGAGACAGCTGGTGGAAACCCGGCAGATCACAAAGCACGTG GCACAGATCCTGGACTCCCGGATGAACACTAAGTACGACGAGAATGACAAGCTGATCCGGGAAGT GAAAGTGATCACCCTGAAGTCCAAGCTGGTGTCCGATTTCCGGAAGGATTTCCAGTTTTACAAAGT GCGCGAGATCAACAACTACCACCACGCCCACGACGCCTACCTGAACGCCGTCGTGGGAACCGCCC TGATCAAAAAGTACCCTAAGCTGGAAAGCGAGTTCGTGTACGGCGACTACAAGGTGTACGACGTG CGGAAGATGATCGCCAAGAGCGAGCAGGAAATCGGCAAGGCTACCGCCAAGTACTTCTTCTACAG CAACATCATGAACTTTTTCAAGACCGAGATTACCCTGGCCAACGGCGAGATCCGGAAGCGGCCTC TGATCGAGACAAACGGCGAAACCGGGGAGATCGTGTGGGATAAGGGCCGGGATTTTGCCACCGTG CGGAAAGTGCTGAGCATGCCCCAAGTGAATATCGTGAAAAAGACCGAGGTGCAGACAGGCGGCT TCAGCAAAGAGTCTATCCTGCCCAAGAGGAACAGCGATAAGCTGATCGCCAGAAAGAAGGACTG GGACCCTAAGAAGTACGGCGGCTTCGACAGCCCCACCGTGGCCTATTCTGTGCTGGTGGTGGCCA AAGTGGAAAAGGGCAAGTCCAAGAAACTGAAGAGTGTGAAAGAGCTGCTGGGGATCACCATCAT GGAAAGAAGCAGCTTCGAGAAGAATCCCATCGACTTTCTGGAAGCCAAGGGCTACAAAGAAGTG AAAAAGGACCTGATCATCAAGCTGCCTAAGTACTCCCTGTTCGAGCTGGAAAACGGCCGGAAGAG AATGCTGGCCTCTGCCGGCGAACTGCAGAAGGGAAACGAACTGGCCCTGCCCTCCAAATATGTGA ACTTCCTGTACCTGGCCAGCCACTATGAGAAGCTGAAGGGCTCCCCCGAGGATAATGAGCAGAAA CAGCTGTTTGTGGAACAGCACAAGCACTACCTGGACGAGATCATCGAGCAGATCAGCGAGTTCTC CAAGAGAGTGATCCTGGCCGACGCTAATCTGGACAAAGTGCTGTCCGCCTACAACAAGCACCGGG ATAAGCCCATCAGAGAGCAGGCCGAGAATATCATCCACCTGTTTACCCTGACCAATCTGGGAGCC CCTGCCGCCTTCAAGTACTTTGACACCACCATCGACCGGAAGAGGTACACCAGCACCAAAGAGGT GCTGGACGCCACCCTGATCCACCAGAGCATCACCGGCCTGTACGAGACACGGATCGACCTGTCTC AGCTGGGAGGCGACAAAAGGCCGGCGGCCACGAAAAAGGCCGGCCAGGCAAAAAAGAAAAAGTAAG

#### Cas9-α2-β2

ATGGACTATAAGGACCACGACGGAGACTACAAGGATCATGATATTGATTACAAAGACGATGACGA TAAGATGGCCCCAAAGAAGAAGCGGAAGGTCGGTATCCACGGAGTCCCAGCAGCCGACAAGAAG TACAGCATCGGCCTGGACATCGGCACCAACTCTGTGGGCTGGGCCGTGATCACCGACGAGTACAA GGTGCCCAGCAAGAAATTCAAGGTGCTGGGCAACACCGACCGGCACAGCATCAAGAAGAACCTG ATCGGAGCCCTGCTGTTCGACAGCGGCGAAACAGCCGAGGCCACCCGGCTGAAGAGAACCGCCA GAAGAAGATACACCAGACGGAAGAACCGGATCTGCTATCTGCAAGAGATCTTCAGCAACGAGAT GGCCAAGGTGGACGACAGCTTCTTCCACAGACTGGAAGAGTCCTTCCTGGTGGAAGAGGATAAGA AGCACGAGCGGCACCCCATCTTCGGCAACATCGTGGACGAGGTGGCCTACCACGAGAAGTACCCC ACCATCTACCACCTGAGAAAGAAACTGGTGGACAGCACCGACAAGGCCGACCTGCGGCTGATCTA TCTGGCCCTGGCCCACATGATCAAGTTCCGGGGCCACTTCCTGATCGAGGGCGACCTGAACCCCG ACAACAGCGACGTGGACAAGCTGTTCATCCAGCTGGTGCAGACCTACAACCAGCTGTTCGAGGAA AACCCCATCAACGCCAGCGGCGTGGACGCCAAGGCCATCCTGTCTGCCAGACTGAGCAAGAGCAG ACGGCTGGAAAATCTGATCGCCCAGCTGCCCGGCGAGAAGAAGAATGGCCTGTTCGGAAACGGTA TTGCCCTGAGCCTGGGCCTGACCCCCAACTTCAAGAGCAACTTCGACCTGGCCGAGGATGCCAAA CTGCAGCTGAGCAAGGACACCTACGACGACGACCTGGACAACCTGCTGGCCCAGATCGGCGACCA GTACGCCGACCTGTTTCTGGCCGCCAAGAACCTGTCCGACGCCATCCTGCTGAGCGACATCCTGAG AGTGAACACCGAGATCACCAAGGCCCCCCTGAGCGCCTCTATGATCAAGAGATACGACGAGCACC ACCAGGACCTGACCCTGCTGAAAGCTCTCGTGCGGCAGCAGCTGCCTGAGAAGTACAAAGAGATT TTCTTCGACCAGAGCAAGAACGGCTACGCCGGCTACATTGACGGCGGAGCCAGCCAGGAAGAGTT CTACAAGTTCATCAAGCCCATCCTGGAAAAGATGGACGGCACCGAGGAACTGCTCGTGAAGCTGA ACAGAGAGGACCTGCTGCGGAAGCAGCGGACCTTCGACAACGGCAGCATCCCCCACCAGATCCAC CTGGGAGAGCTGCACGCCATTCTGCGGCGGCAGGAAGATTTTTACCCATTCCTGAAGGACAACCG GGAAAAGATCGAGAAGATCCTGACCTTCCGCATCCCCTACTACGTGGGCCCTCTGGCCAGGGGAA ACAGCAGATTCGCCTGGATGACCAGAAAGAGCGAGGAAACCATCACCCCCTGGAACTTCGAGGA AGTGGTGGACAAGGGCGCTTCCGCCCAGAGCTTCATCGAGCGGATGACCAACTTCGATAAGAACC TGCCCAACGAGAAGGTGCTGCCCAAGCACAGCCTGCTGTACGAGTACTTCACCGTGTATAACGAG CTGACCAAAGTGAAATACGTGACCGAGGGAATGAGAAAGCCCGCCTTCCTGAGCGGCGAGCAGA AAAAGGCCATCGTGGACCTGCTGTTCAAGACCAACCGGAAAGTGACCGTGAAGCAGCTGAAAGA GGACTACTTCAAGAAAATCGAGTGCTTCGACTCCGTGGAAATCTCCGGCGTGGAAGATCGGTTCA ACGCCTCCCTGGGCACATACCACGATCTGCTGAAAATTATCAAGGACAAGGACTTCCTGGACAAT GAGGAAAACGAGGACATTGGTGAAGATATCGTGCTGACCCTGACACTGTTTGAGGACAGAGAGAT GATCGAGGAACGGCTGAAAACCTATGCCCACCTGTTCGACGACAAAGTGATGAAGCAGCTGAAGC GGCGGAGATACACCGGCTGGGGCAGGCTGAGCCGGAAGCTGATCAACGGCATCCGGGACAAGCA GTCCGGCAAGACAATCCTGGATTTCCTGAAGTCCGACGGCTTCGCCAACAGAAACTTCATGCAGC TGATCCACGACGACAGCCTGACCTTTAAAGAGGACATCCAGAAAGCCCAGGTGTCCGGCCAGGGC GATAGCCTGCACGAGCACATTGCCAATCTGGCCGGCAGCCCCGCCATTAAGAAGGGCATCCTGCA GACAGTGAAGGTGGTGGACGAGCTCGTGAAAGTGATGGGCCGGCACAAGCCCGAGAACATCGTG ATCGAAATGGCCAGAGAGAACCAGACCACCCAGAAGGGACAGAAGAACAGCCGCGAGAGAATG AAGCGGATCGAAGAGGGCATCAAAGAGCTGGGCAGCCAGATCCTGAAAGAACACCCCGTGGAAA ACACCCAGCTGCAGAACGAGAAGCTGTACCTGTACTACCTGCAGAATGGGCGGGATATGTACGTG GACCAGGAACTGGACATCAACCGGCTGTCCGACTACGATGTGGACCATATCGTGCCTCAGAGCTT TCTGAAGGACGACTCCATCGACAACAAGGTGCTGACCAGAAGCGACAAGAACCGGGGCAAGAGC GACAACGTGCCCTCCGAAGAGGTCGTGAAGAAGATGAAGAACTACTGGCGGCAGCTGCTGAACG CCAAGCTGATTACCCAGAGAAAGTTCGACAATCTGACCAAGGCCGAGAGAGGCGGCCTGAGCGA ACTGGATAAGGCCGGCTTCATCAAGAGACAGCTGGTGGAAACCCGGCAGATCACAAAGCACGTG GCACAGATCCTGGACTCCCGGATGAACACTAAGTACGACGAGAATGACAAGCTGATCCGGGAAGT GAAAGTGATCACCCTGAAGTCCAAGCTGGTGTCCGATTTCCGGAAGGATTTCCAGTTTTACAAAGT GCGCGAGATCAACAACTACCACCACGCCCACGACGCCTACCTGAACGCCGTCGTGGGAACCGCCC TGATCAAAAAGTACCCTAAGCTGGAAAGCGAGTTCGTGTACGGCGACTACAAGGTGTACGACGTG CGGAAGATGATCGCCAAGAGCGAGCAGGAAATCGGCAAGGCTACCGCCAAGTACTTCTTCTACAG CAACATCATGAACTTTTTCAAGACCGAGATTACCCTGGCCAACGGCGAGATCCGGAAGCGGCCTC TGATCGAGACAAACGGCGAAACCGGGGAGATCGTGTGGGATAAGGGCCGGGATTTTGCCACCGTG CGGAAAGTGCTGAGCATGCCCCAAGTGAATATCGTGAAAAAGACCGAGGTGCAGACAGGCGGCT TCAGCAAAGAGTCTATCCTGCCCAAGAGGAACAGCGATAAGCTGATCGCCAGAAAGAAGGACTG GGACCCTAAGAAGTACGGCGGCTTCGACAGCCCCACCGTGGCCTATTCTGTGCTGGTGGTGGCCA AAGTGGAAAAGGGCAAGTCCAAGAAACTGAAGAGTGTGAAAGAGCTGCTGGGGATCACCATCAT GGAAAGAAGCAGCTTCGAGAAGAATCCCATCGACTTTCTGGAAGCCAAGGGCTACAAAGAAGTG AAAAAGGACCTGATCATCAAGCTGCCTAAGTACTCCCTGTTCGAGCTGGAAAACGGCCGGAAGAG AATGCTGGCCTCTGCCGGCGAACTGCAGAAGGGAAACGAACTGGCCCTGCCCTCCAAATATGTGA ACTTCCTGTACCTGGCCAGCCACTATGAGAAGCTGAAGGGCTCCCCCGAGGATAATGAGCAGAAA CAGCTGTTTGTGGAACAGCACAAGCACTACCTGGACGAGATCATCGAGCAGATCAGCGAGTTCTC CAAGAGAGTGATCCTGGCCGACGCTAATCTGGACAAAGTGCTGTCCGCCTACAACAAGCACCGGG ATAAGCCCATCAGAGAGCAGGCCGAGAATATCATCCACCTGTTTACCCTGACCAATCTGGGAGCC CCTGCCGCCTTCAAGTACTTTGACACCACCATCGACCGGAAGAGGTACACCAGCACCAAAGAGGT GCTGGACGCCACCCTGATCCACCAGAGCATCACCGGCCTGTACGAGACACGGATCGACCTGTCTC AGCTGGGAGGCGACAAAAGGCCGGCGGCCACGAAAAAGGCCGGCCAGGCAAAAAAGAAAAAGT

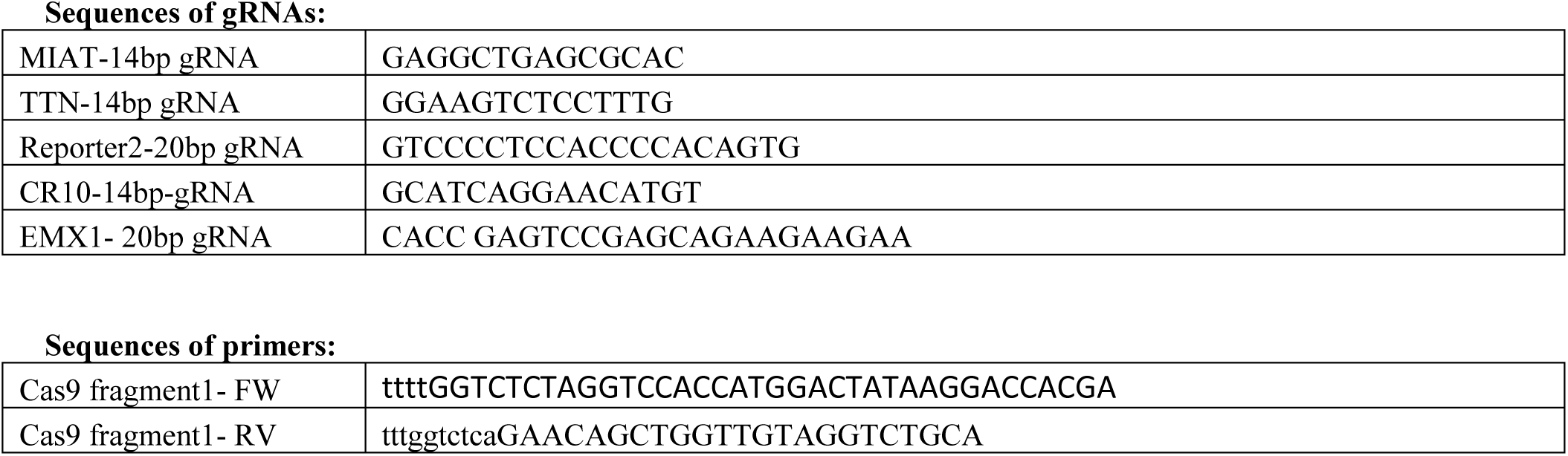
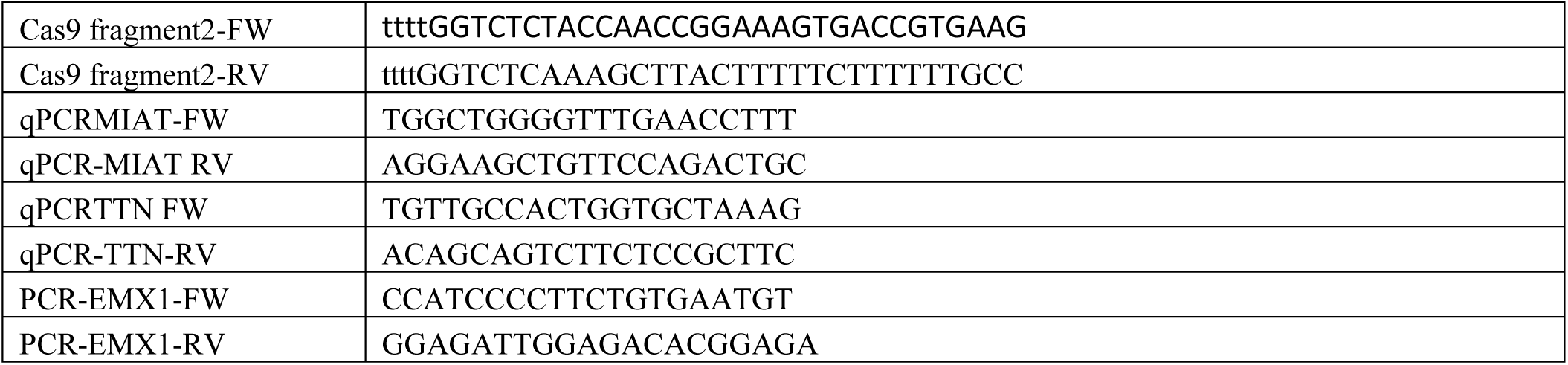

